# The endoribonuclease Rae1 from *Bacillus subtilis* cleaves mRNA upstream of stalled ribosomes

**DOI:** 10.1101/2025.06.04.657793

**Authors:** Valentin Deves, Alexandre D’Halluin, Laëtitia Gilet, Ciarán Condon, Frédérique Braun

**Author notes:** These authors contributed equally and are listed alphabetically (Valentin Deves, Alexandre D’Halluin).

## Abstract

The ribosome-associated endoribonuclease 1 (Rae1) cleaves mRNAs in a translation-dependent manner. Here, we identify a new Rae1 target, the *fliY* mRNA, which is cleaved by Rae1 in the absence of the elongation factor P (EF-P). The Rae1 site was mapped 12 nucleotides upstream of the second proline codon of an SPP stalling motif in *fliY*. Remarkably, Rae1 cleavages also occur 12 nucleotides upstream of the stop codon within two validated Rae1 mRNA targets, *bmrX* and *spyA* (*S1025*). Shifting the stop codon relative to the Rae1 cutting site abolished Rae1 sensitivity of *bmrX* and *spyA* mRNAs. We show that ribosome pausing occurs at the *spyA* stop codon, confirming its crucial role, and positioning the Rae1 cleavage at the tail end of the stalled ribosome, rather than in the A-site as previously proposed. These findings reveal a compelling novel mechanism by which Rae1 mediates mRNA cleavage in coordination with immobile ribosomes.

## Introduction

Translation and mRNA decay are closely interdependent, ensuring that mRNAs are degraded when they are no longer needed for protein synthesis ^1–4^. Translation efficiency, governed by initiation rates and codon bias, is known to directly affect mRNA stability, where inefficiently translated mRNAs are more likely to be targeted for decay ^5,6^. In bacteria, ribosomes can impact mRNA degradation either by blocking 5’-exoribonucleases ^7–9^ or by hindering access to endoribonucleolytic cleavage sites ^10–13^. Transcription and translation are tightly coupled in *Escherichia coli* ^14^, efficiently limiting access to nascent transcripts by RNases, while these two processes were recently shown to be mostly disconnected in *Bacillus subtilis* ^15^. This, added to the fact that the RNA degradation machineries of these two organisms are very different^16^, underscores the potential divergence in the interplay between translation and RNA decay between Gram-negative and Gram-positive bacteria.

The translation machinery is also a key player in surveillance systems for mRNA quality control ^17–21^. In yeast, the Cue2 RNase was identified as an effector of the No-Go decay (NGD) pathway that initiates the degradation of defective mRNAs on which ribosomes collide and contributes to the rescue of stalled ribosomes ^22^. Only a few bacterial RNases have been described to interact with the ribosome and cleave mRNAs in a translation-dependent manner ^23–26^. Among them, SmrB, a homolog of yeast Cue2, was recently shown to cut mRNAs between stalled and collided ribosomes, triggering ribosome rescue via the *trans*-translation pathway in *E. coli* ^19,27^. Strikingly, its *B. subtilis* homolog, MutS2, was proposed to rescue collided ribosomes by splitting them into individual subunits via the ribosome quality control (RQC) pathway without promoting RNA cleavage ^28,29^. We have shown that the ribosome associated endoribonuclease 1 (Rae1) from *B. subtilis* cleaves mRNA in a translation-dependent manner and promotes ribosome rescue via the *trans*-translation pathway involving the transfer-messenger RNA (tmRNA) ^26,30^. It has no homolog in *E. coli* but it is widely conserved among Firmicutes, Cyanobacteria and the chloroplasts of higher plants ^31^. Rae1 shares no structural homology with MutS2, and we have shown that Rae1 and MutS2 act in distinct pathways ^30^.

The two Rae1 cleavage sites characterized thus far were mapped within short open reading frames (ORFs), *spyA* and *bmrX,* encoding 17-aa and 26-aa peptides, respectively ^26,30^. SpyA (S1025), which belongs to the *spyTA* polycistronic mRNA, was recently shown to act as an antidote to the SpyT (YrzI) toxin ^32^. BmrX, encoded by the *bmrBXCD* operon, is a newly-discovered cryptic ORF located between the antibiotic-sensitive attenuator *bmrB* and *bmrCD,* encoding a heterodimeric ABC multidrug exporter ^30,33,34^. Cleavage by Rae1 shuts off leaky readthrough expression of *bmrCD* in the absence of antibiotics. Both Rae1 cleavage sites are located within RNA sequences that encode well-conserved amino acids in the Bacillales, EKIEGG for BrmX and MEKDQV for SpyA. In both cases, Rae1 requires translation of the correct reading frame to function ^26,30^. However, the nucleotide and the amino acid sequences of these two Rae1 targets do not share any common features, raising the question of what constitutes an mRNA target for Rae1.

Here, we identify a new Rae1 site within the *fliY* mRNA, whose cleavage by Rae1 depends on an SPP motif at which the ribosome stalls in the absence of elongation factor P (EF-P). We show that ribosomes also pause on the *spyA* mRNA with the stop codon in its A-site. We also show that the distance between the Rae1 cleavage site and the stop codon of *bmrX* and *spyA* is crucial for Rae1 cleavage, and we identify new sequence elements within Rae1 mRNA targets required to trigger Rae1 cleavage. This study points to a novel mechanism of mRNA quality control involving ribosome pausing and mRNA cleavage by the endoribonuclease Rae1.

## Materials and Methods

### Strains and constructs

Oligonucleotides and strains used in this study are shown in Supplemental Tables S1 and S2 respectively. The *B. subtilis* strains used in this study were derivatives of W168 (lab strain SSB1002).

The pHM2-*bmrX-gfp* (pl904) plasmid containing the *spyA* Shine-Dalgarno sequence followed by the *bmrX* ORF translationally fused upstream of GFP was obtained by PCR amplification using oligo pair CC2888/CC572 using pPspac-*bmrBX-gfp f0* (pl845) plasmid as a template ^30^. The pHM2-*spyA(+18 nts)* (pl973) and pHM2-*spyA UAG(+18 nts)* (pl974) plasmids were constructed by PCR amplification using oligo pair CC2962/CC572, with pHM2-*hbs*Δ*-spyA(+18 nts)* (pl895) or pHM2-*hbs*Δ*-spyA UAG(+18 nts)* (pl927) plasmids as a template. The pHM2-*fliY* plasmid (pl856) was constructed by PCR amplification using oligo pair CC2696/CC2697, with chromosomal DNA from SSB1002 as a template. The pHM2-*fliY*Δ (pl922) plasmid containing a truncated version of the *fliY* mRNA was constructed by PCR amplification using oligo pair CC2696/CC2950 with pHM2-*fliY* (pl856) as a template. The construction of the additional plasmids is fully described in Supplemental Table S3. All of the inserts listed in Table S3 were obtained by overlapping PCR. They were subcloned between the HindIII and BamHI site of the integrative pHM2 plasmid and placed under the control of a constitutive promoter (Pspac(con)). All plasmid constructs were verified by sequencing.

All derivatives of the pHM2 integrative plasmid were linearized with XbaI, before transformation, for integration of the constructs into the *amyE* locus of strains SSB1002 (WT), CCB375 *(Δrae1*), CCB1481 (*Δefp*) or CCB1482 (*Δrae1Δefp*) (Table S2). The CCB375 strain *(Δrae1*) is resistant to erythromycin. The CCB1481 and CCB1482 strains (*Δefp* and *Δrae1Δefp*) are resistant to kanamycin and to erythromycin and kanamycin, respectively. The pHM2 vectors confer chloramphenicol resistance.

Strains CCB1479, CCB1480, CCB1481 and CCB1482 were constructed by transferring the *efp::kan* construct from strain BKK24450 in the single-gene deletion library ^59^ into CCB1456 (pHM2*-fliY*Δ), CCB1461 (*Δrae1* pHM2*-fliY*Δ), SSB1002 (WT) and CCB375 *(Δrae1*), respectively. Strains CCB1588 and CCB1596 were constructed by transferring the *rnjA::spec* construct from strain CCB434 into CCB1479 (*Δefp* pHM2*-fliY*Δ) and CCB1480 (*Δrae1Δefp* pHM2*-fliY*Δ).

The pNAR913 (*Bs_prfA-his6*) and pNAR915 (*Bs_prfB-his6*) plasmids, overexpressing hexahistidine-tagged RF1 and RF2 from the pET28b-based plasmid were a gift from Shinobu Chiba ^60^. These plasmids were transformed into BL21 CodonPlus RIL cells to yield strain CCE300 and CCE299, respectively.

### RNA isolation and Northern blots

Northern blots were performed on total RNA isolated from *B. subtilis* cells growing in 2xYT medium either by the glass beads/phenol method described in ^61^ or by the RNA-Snap method described in ^62^. Northern blots were performed as described previously ^63^. Northern blots were exposed to PhosphorImager screens (GE Healthcare), and the signal was obtained by scanning with a Typhoon scanner (GE Healthcare) and analyzed by Fiji (ImageJ) software.

### Primer extension assays

Primer extension assays were performed on total RNA extracted from *B. subtilis* cells growing in 2xYT medium by the phenol method as described previously ^64^. 20 μg (qsp 8.5 μl in H_2_0) of RNA were denatured 3 min at 95°C, immediately followed by a 3 min incubation at 4°C on ice. Then, 1 μl (0.6 pmol) of 5’-labelled (^32^P) probe was added, and reactions were incubated at 65°C for 5 min, followed by a 10 min incubation at 35°C. A 5.5 μl mix containing 10 mM each dNTP and 200 units M-MLV reverse transcriptase (Invitrogen) in RT M-MLV buffer was added to the reactions for a final volume of 15 μl, before incubation at 35°C for 2 min, followed by a 45 min incubation at 42°C and a 5 min incubation at 55°C. Reactions were stopped with 5 μl RNA loading dye (Biolabs) and 10 μl were loaded on 5% sequencing gels. The DNA template was sequenced using the CC2947 primer and the Thermo Sequenase Cycle Sequencing Kit (Applied Biosystems). Oligo CC2947 was used as the probe to map the Rae1 cleavage site within the *fliY*Δ ORF.

### Protein production and purification

The Rae1, RF1 and RF2 hexahistidine-tagged proteins from *B.subtilis* were produced in *E. coli* using the plasmids : pET28-Rae1Chis ^26^, pET28-RF1His6 (pNAR913) and pET28-RF2His6 (pNAR915) ^60^. Rae1, RF1 and RF2 expression was induced with 1 mM IPTG for 4 h and purified on Ni-NTA resin according to previously published protocols ^26^. Peak fractions were dialyzed against 20 mM Tris pH 8.0, 150 mM NaCl, 10% glycerol, concentrated using Amicon filters with 3-kDa cut-off and applied to a Superdex 26/60 sizing column equilibrated in the same buffer. Peak fractions were reconcentrated for *in vitro* trials.

### *In vitro* translation of SpyA peptides

*In vitro* translation reactions were typically performed using the PURExpress Δribosomes kit (New England Biolabs) in 5 μl reactions with 2.5 pmol of *B. subtilis* ribosomes and with [35-S]-L-methionine using 100 ng of PCR template to transcribe *spyA* mRNA by T7 RNA polymerase. Reactions were incubated at 37°C for 30 minutes and precipitated with 500 µl TCA 5%. The pellets were washed with acetone and resuspended in 12 µl buffer (10 mM Tris pH 7.5, 100 mM NaCl). The reactions were run on 20% SDS-PAGE. The gel was dried for 90 min and revealed using the Typhoon FLA 9500 5GE Healthcare Life Sciences).

### *In vitro* translation Rae1 cleavage assay

70S ribosomes were purified from 1 g of frozen *B. subtilis* W168 cells as previously described ^26^. *spyA* mRNA mutants were transcribed by T7 RNA polymerase *in vitro* using a MegaShortScript kit (Ambion) and PCR fragments as templates. Templates were amplified from the indicated plasmid using oligo pairs CC1660/CC429 (*spyA* pl707*, spyA(+18nts)* pl895 and *spyA UAG(+18nts)* pl927 and *spyA(14-*△*LRM)* pl937) or CC1660/CC1661 (*spyA* pl925*, spyA-AAmod* pl1008 and *spyA(UGA)* pl941). The *in vitro* transcripts were then purified as previously described (Leroy et al., 2017). *In vitro* translation assays (PURExpress ribosomes; New England Biolabs) were typically performed in 5 μl reactions with 1 pmol 5^’^-labelled (^32^P) mRNA and 6.5 pmol *B. subtilis* 70S ribosomes. Rae1 was added at 38 pmol per reaction and RF1 and RF2 at 2 or 20 pmol per reaction. Reactions were incubated at 37°C for 30 min followed by phenol extraction. The aqueous phase was precipitated overnight at −20°C, or 2h at −80°C, in 3 volumes of ethanol and 1/10 vol of LiCl, supplemented with 1 μl glycogen (20 mg/ml). Pellets were resuspended in 6 μl H_2_O, with 6 μl RNA loading dye (Biolabs). 6 μl of the samples were run on 5% sequencing gels. The gel was dried for 90 min and revealed using the Typhoon FLA 9500 5GE Healthcare Life Sciences).

### Toeprint assay

The position of stalled ribosomes on mRNA was determined by toeprinting assay. RNAs were prepared as for the *in vitro* assays of Rae1 activity. A total of 1 pmol of RNA template was used for *in vitro* translation with 3.3 µM (2 pmol) of 70S ribosomes from SSB1002 strain and using the PURExpress Δribosome kit (New England Biolabs) following the manufacturer’s recommendations. When necessary, reactions were incubated with either tetracycline or puromycin (100 µg/ml)) to enhance ribosome stalling. The *in vitro* translation reactions were incubated at 37°C for 30 min and then were directly incubated for 5 min at 37°C with 1 pmol of radiolabeled probe CC1661 followed by incubation 5 min on ice and 2 min at room temperature. The DNA probe was 5’ radiolabeled using ^32^P-γ ATP and PNK enzyme (New England Biolabs) for 45 min at 37°C followed by a G50 column purification (Cytiva). RNAs were reverse transcribed using 0.5 µl of AMV Reverse Transcriptase (Promega), 0.1 mM dNTP and 1X of Pure System Buffer (5 mM K-phosphate pH 7.3, 9 mM MgOAc, 95 mM K-glutamate, 5LmM NH_4_Cl, 0.5LmM CaCl_2_, 1 mM spermidine, 8 mM putrescine, 1 mM DTT). Reactions were incubated 20 min at 37°C and directly purified by a phenol purification. Pellets were dried for 30 sec using a vacuum dryer and resuspended in 5 µl of water and 5 µl of 95% formamide loading dye. Reactions were run on 5% sequencing gels. The DNA template was sequenced using the CC1661 primer and the Thermo Sequenase Cycle Sequencing Kit (Applied Biosystems). The gel was dried for 90 min and revealed using the Typhoon FLA 9500 5GE Healthcare Life Sciences).

## Results

### Rae1 cleavage within the *fliY* mRNA is promoted by ribosome pausing at the SPP motif

A previous report showed that a swarming defect in strains lacking the *efp* gene, encoding the translation elongation factor P (EF-P), can be genetically suppressed by nonsense mutations in the *rae1* gene ^35^. EF-P binds to the E-site of the ribosome and accelerates peptide bond formation between consecutive prolines, thereby preventing ribosome stalling at these difficult-to-translate sequences. It was shown that in the Δ*efp* strain, ribosomes pause at an SPP motif, corresponding to codons 164–166 of the flagellar FliY ORF and that the swarming defect results from a strong decrease in expression of the *fliY* mRNA. We hypothesized that suppression of the swarming defect in the Δ*rae1* <u>Δ</u>*efp* strain might be due to the stabilization of the *fliY* mRNA, allowing sufficient synthesis of FliY to make functional flagella.

The *fliY* gene belongs to a 32-gene polycistronic operon whose transcript is difficult to analyze by Northern blot. To determine whether Rae1 impacts the stability of the *fliY* mRNA, we constructed a truncated version of the *fliY* gene (*fliY*Δ) encoding the first 180 amino acids (including the ^164^SPP^166^ motif) with its native 5’ and 3’ UTRs, inserted at the *amyE* locus. The stability of the *fliY*Δ mRNA was analyzed by Northern blot in WT, Δ*rae1*, Δ*efp* and Δ*efp* Δ*rae1* strains. The *rae1* deletion had no impact on *fliY*Δ mRNA stability in the presence of EF-P (Fig 1A). In contrast, the *fliY*Δ transcript was stabilized 2.8-fold in the Δ*efp* Δ*rae1* double mutant strain compared to the Δ*efp* mutant strain (6.8 and 2.5 min half-lives, respectively), indicating that Rae1 promotes the degradation of the *fliY*Δ transcript in the absence of EF-P (Fig. 1A). We mutated the two CCU proline codons 165–166 to alanine codons (GCU) to test whether the SPP motif was required for Rae1 cleavage. The *fliY*Δ *(P165A P166A)* transcript was less sensitive to Rae1 (1.3-fold stabilization in the Δ*efp rae1* strain), suggesting that the pausing of the ribosome at the proline dimer promotes Rae1 cleavage (Fig. 1A).

**Figure 1:**
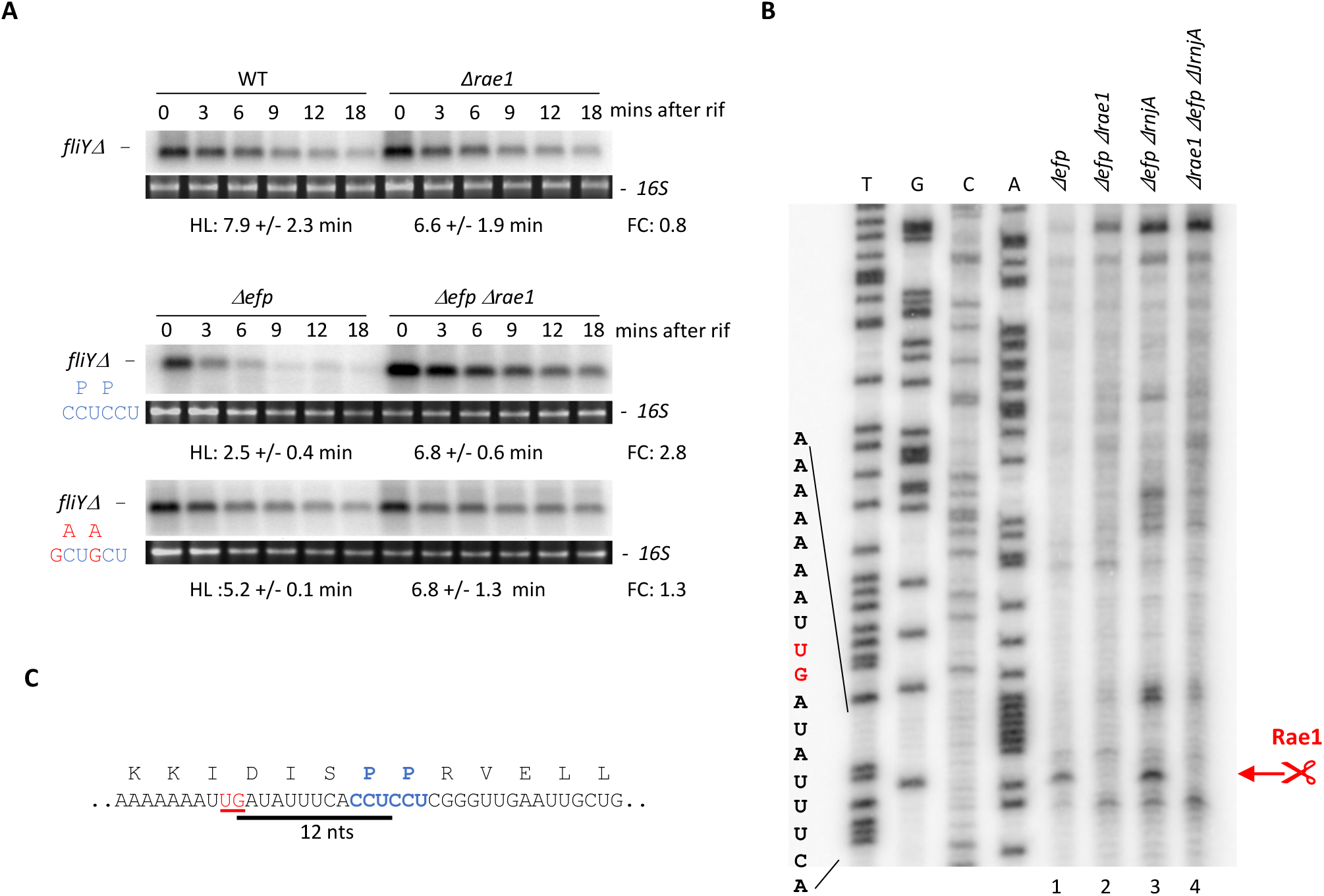
Rae1 cleaves the *fliY* mRNA upstream of the SPP stalling motif. A. Stability of *fliY* mRNA revealed by Northern Blot. The stability of the truncated version of *fliY (fliY*△) and its derivative with the SPP motif mutated to SAA (*fliY*△ *(P165A P166A)),* inserted into the chromosome at the *amyE* locus, was analyzed in WT, △*rae1,* △*efp* and △*efp*△*rae1* strains. RNAs were extracted 0, 3, 6, 9, 12, and 18 min after addition of rifampicin at 150 µg/ml. The blots were probed with oligonucleotide CC1661. Half-lives (HL) were calculated from biological replicates. The fold-change (FC) in half-lives is given next to each autoradiogram. B. Cleavage site of *fliY*△ mRNA by Rae1 determined by primer extension assay. Total RNA from △*efp,* △*efp*△*rae1,* △*efp*△*rnjA* and △*efp*△*rae1*△*rnjA* strains containing the *fliY*Δ gene inserted at *amyE* were subject to reverse transcription. Primer extension was performed with oligonucleotide CC1661. The Rae1 cleavage site is highlighted in red. C. Localization of Rae1 cleavage site on the *fliY* mRNA. The Rae1 cleavage site (underlined in red) is located 12 nts upstream of the second proline codon of the SPP stalling motif (in bold). The 12-nt distance between the proline codon and the Rae1 cleavage site is represented by a black line.

We then mapped the Rae1 cleavage site by performing primer extension assays on total RNA isolated from Δ*efp* or Δ*efp* Δ*rae1* strains containing the *fliY*Δ gene in the *amyE* locus. This experiment was also done in a Δ*rnjA* background to protect the downstream cleavage product from 5’-3’ degradation, if necessary. In the two strains lacking EF-P (Δ*efp* and Δ*efp* Δ*rnjA*), we detected a reverse transcription stop that mapped 12 nts upstream of the proline codon at position 166, which was not seen in the absence of Rae1, suggesting that this is the site of Rae1 cleavage (Fig. 1B and 1C). Since a ribosome footprint ranges from 11-15 nt on either side of the A-site, depending on whether the ribosome is collided, elongating or initiating, this would suggest that Rae1 cleaves the *fliY*Δ mRNA at the tail end of the stalled ribosome, with the second proline codon in its A-site.

Our results show that the *fliY* mRNA is a new target of the Rae1 and that ribosome pausing at the ^164^SPP^166^ motif promotes Rae1 cleavage. Ribosome pausing at the SPP motif in the absence of EF-P would lead to irreversible cleavage of the *fliY* mRNA by Rae1. However, in the absence of this RNase, ribosomes would have enough time to resolve the translation impediment of the *fliY* mRNA, leading to the suppression of the swarming defect.

### Ribosome pausing occurs at the *spyA* stop codon

Based on the evidence that ribosome pausing plays a role in Rae1 cleavage of the *fliY* mRNA, we investigated whether ribosome pausing also occurred on the *spyA* mRNA. We performed toeprint experiments on *in vitro* translation reactions using the WT *spyA* transcript as a template and an oligonucleotide hybridizing to the 3’-UTR for primer extension. As negative controls, we performed reactions without ribosomes, or in the presence of the translation inhibitors tetracycline or puromycin. Tetracycline inhibits protein synthesis by binding to the A-site of the ribosome, which prevents aminoacyl-tRNA entry. Puromycin is a structural analog of aminoacyl tRNAs that enters the ribosomal A-site and is incorporated into the newly synthesized protein, leading to spontaneous dissociation of the nascent peptide from the ribosome. Stalled ribosomes do not actively incorporate puromycin and are therefore resistance to its effects ^36^. As expected, a toeprint signal was observed at position +17 relative to the A of the start codon, corresponding to the ribosomal initiation complex, in all reactions containing ribosomes (Fig. 2, lanes 2-4). A second toeprint signal was observed at position +17 relative to the U of the stop codon, consistent with a stalled ribosome with the stop codon in its A-site (Fig. 2, lane 2). In agreement, this toeprint signal codon was still detected in the presence of puromycin (Fig. 2, lane 4) while it is was not observed in the negative control sample without ribosomes or in the presence of tetracycline (Fig. 2, lanes 1 and 3). These data confirm that ribosomes stall on the *spyA* mRNA stop codon.

**Figure 2:**
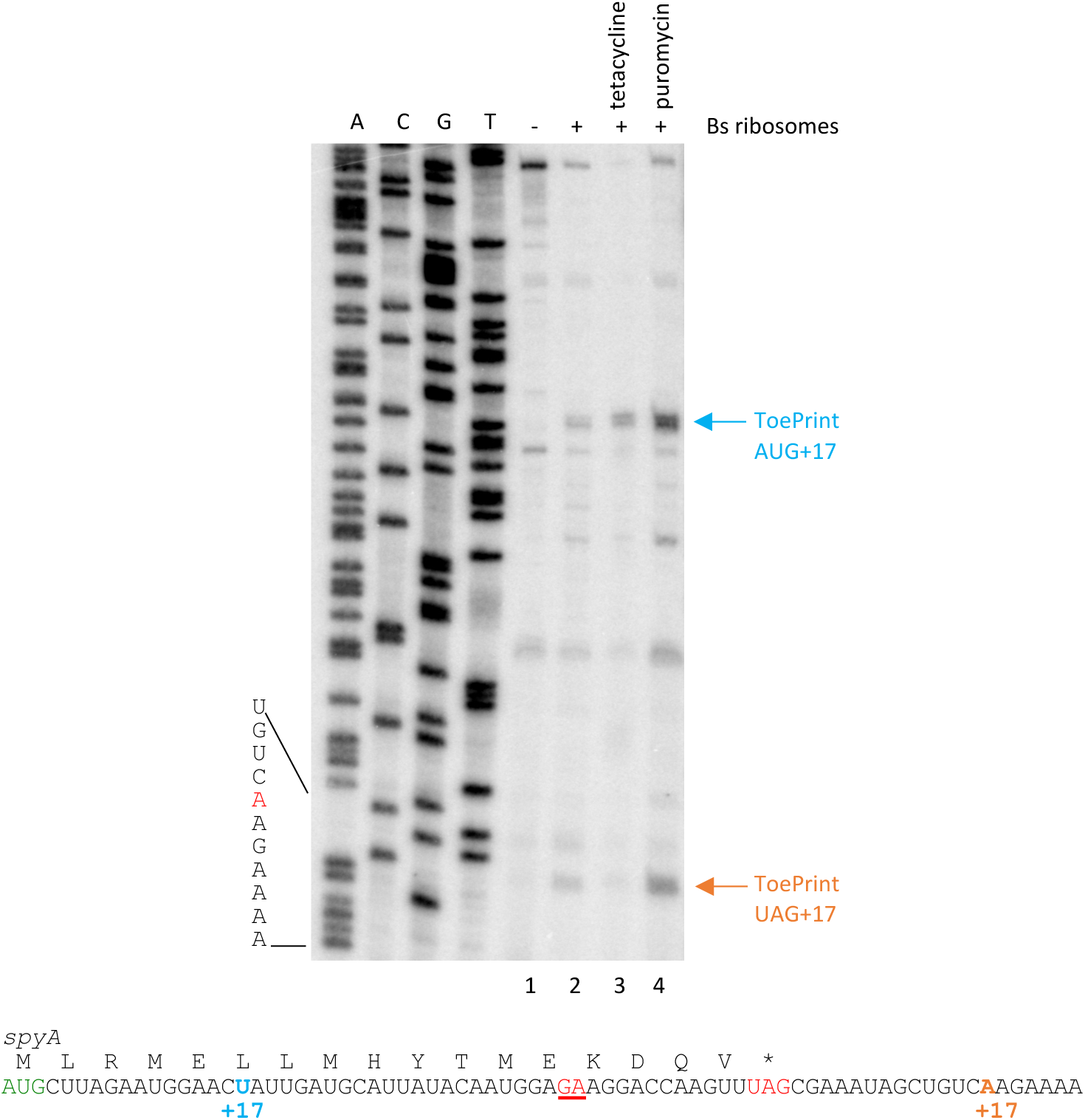
Ribosomes stall at the *spyA* stop codon. Ribosome pausing on *spyA* revealed by toe-print assay. The *in vitro-*transcribed *spyA* mRNA was translated *in vitro* and toe-print assays performed by primer extension using primer CC1661. RNAs were incubated without (-) (lane 1) or with 2.7 pmol of *B. subtilis* ribosomes (+) (lanes 2-4), in the presence of 100 µg/ml of tetracycline (lane 3) or 100 µg/ml of puromycin (lane 4). Blue arrow: toe-print at +17 from start site. Orange arrow: toe-print at +17 from termination site. The sequence of *spyA* is shown in the lower panel. Green: start codon. Red: stop codon. Blue and orange: toeprint signals. Red underlined: Rae1 cleavage site.

### The distance between the Rae1 site and the stop codon is important for Rae1 cleavage

Although they have no common sequence features, we noticed that the Rae1 cleavage sites of all three known substrates, *fliY*, *spyA* and *bmrX,* are located 12 nts upstream of a known or potential ribosome pause site, a proline codon in the case of *fliY* and a stop codon for both *spyA* and *bmrX* (Fig 3A) ^26,30^. To determine whether this distance was important for Rae1 cleavage, we moved the *spyA* stop codon 30 nts downstream of the Rae1 site by introducing a random 18-nt sequence after the last sense codon, giving rise to the *spyA (+18 nts)* construct (Fig. 3A). The wild-type (WT) *spyA* and *spyA (+18 nts)* transcripts were expressed from the *amyE* locus, and contained 33 nts upstream of the start codon and 69 nts downstream of the stop codon ^30^. Their stability was measured in WT and Δ*rae1* strains by Northern blot. In contrast to the WT *spyA* transcript, which was stabilized 5.9-fold in the Δ*rae1* strain, the half-life of the *spyA (+18 nts)* mRNA was similar in the WT and Δ*rae1* strain (7.9 min and 7.4 min, respectively) (Fig. 3B). Recreation of a stop codon 12 nts from the Rae1 site in this new context by adding UA upstream of the 18-nt insertion (*spyA UAG(+18 nts)*; Fig 3A), was sufficient to restore Rae1-sensitivity (8.5-fold) (Fig. 3B). Thus, the distance between the Rae1 cleavage site and the stop codon appears important for Rae1 function.

**Figure 3:**
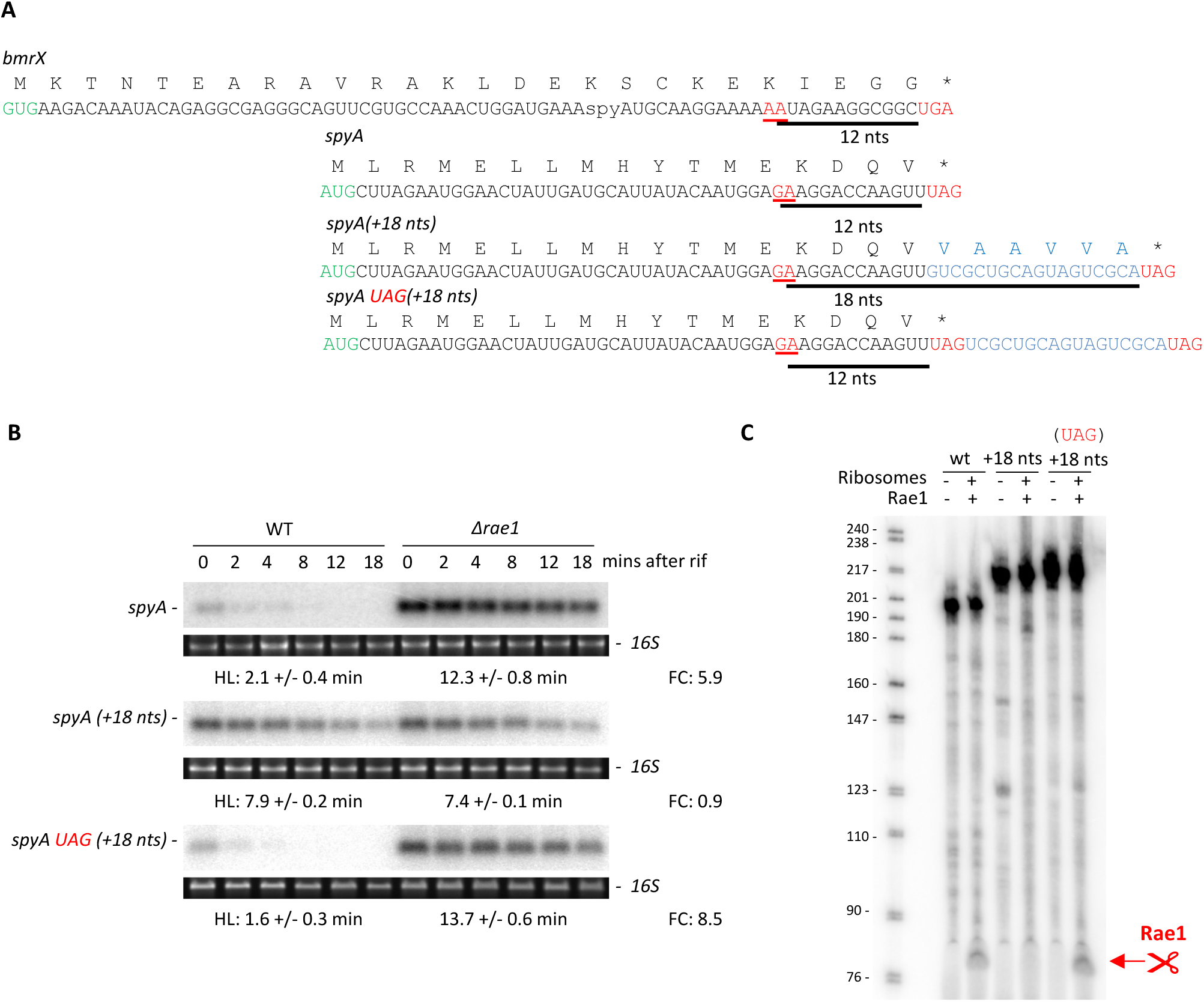
The distance between the Rae1 cleavage site and the stop codon is important for Rae1 cleavage. A. Rae1 cleavage sites in the *bmrX* and *spyA* mRNAs. The Rae1 cleavage site is underlined in red. The nucleotide distance between the stop codon and the Rae1 site is represented by a black line. Start codons are highlighted in green, stop codons in red and the 18-nt random sequence added downstream of *spyA* mRNA in blue (*spyA (+18nts))*. In the *spyA UAG(+18 nts)* construct, a stop codon was reintroduced at its original location. B. Stability of various *spyA* transcripts in WT and △*rae1* strains revealed by Northern blot. Total RNA extracted 0, 2, 4, 8, 12 and 18 min after addition of rifampicin at 150 µg/ml. The blots were probed with oligonucleotide CC2799 that hybridized to *spyA*. The half-lives (HL) of *spyA*, *spyA (+18 nts)* and *spyA UAG(+18 nts*) were calculated from two biological replicates and are given under each autoradiogram. The fold-change (FC) in half-life is given next to each blot C. *In vitro* cleavage assay of *spyA* by Rae1 during translation. 1 pmol of 5’-labelled *spyA, spyA (+18 nts)* and *spyA UAG(+18 nts)* transcripts were incubated in presence of 2.7 pmol of ribosomes and 38 pmol of Rae1. A control for RNA integrity was added without ribosomes or Rae1. The 72-nt band corresponding to the cleaved transcript is highlighted in red.

Rae1 cleavage of these three *spyA* mRNA variants was recapitulated in a purified *in vitro* translation system primed with 70S ribosomes isolated from *B. subtilis*. 5’-labelled *spyA (WT), spyA (+18 nts)* and *spyA UAG(+18 nts)* were *in vitro* transcribed and added to *in vitro* translation assays in the presence or absence of Rae1 (Fig. 3C). A Rae1 cleavage product of the expected size (72 nts) was observed with both the WT *spyA* and the *spyA UAG(+18 nts)* transcripts, showing that Rae1 cleaves the *spyA UAG(+18 nts)* transcript at the same position as WT *spyA,* 12 nts upstream of the stop codon (Fig. 3C). In agreement with the *in vivo* data, the *spyA (+18 nts)* mRNA was not cleaved *in vitro*. These data were further corroborated using a construct in which the *spyA* ORF was inserted in the 3’-UTR of a truncated version of the *hbs* gene (*hbs*Δ*-spyA*), previously shown to be strongly sensitive to Rae1-mediated destabilization ^26^. Consistent with the experiment described above, insertion of *spyA (+18 nts)* in the *hbs*Δ 3’-UTR was no longer able to destabilize *hbs*Δ, whereas insertion of *spyA UAG*(+18 nts) promoted strong Rae1-dependent destabilization, similar to WT *spyA* (Fig. Sup1).

To test whether the distance between the Rae1 cleavage site and the stop codon was also important for Rae1 cleavage within *bmrX*, we took advantage of the *spyA* construct described above, replacing the *spyA* ORF with either the WT *bmrX* ORF or *bmrX* with a different 18-nt random sequence after the last sense codon, *bmrX(+18 nts)* (Fig. Sup2A). A stop codon 12 nts from the Rae1 site was recreated in this new context, by adding UG upstream of the 18-nt insertion, giving rise to *bmrX UGA(+18 nts)* (Fig. Sup2A). Similar to *spyA*, the WT and *bmrX UGA(+18 nts)* mRNAs were both stabilized in the Δ*rae1* strain, 3.6-fold and 4.4-fold respectively, while the *bmrX (+18 nts)* mRNA was insensitive to Rae1 (Fig. Sup2B).

Based on these data, we hypothesized that a *spyA-gfp* mRNA, in which the *spyA* ORF was fused in frame upstream of *gfp* would be insensitive to Rae1 (due to the lack of a stop codon 12 nts downstream of the Rae1 cleavage site), whereas a *gfp-spyA* construct, in which the *spyA* ORF was fused in frame downstream of *gfp* would be Rae1-sensitive (Fig. 4A). Supporting our hypothesis, the half-life of the *spyA-gfp* transcript was independent of Rae1, with half-life values of 12.7 min and 11.7 min in the WT and Δ*rae1* strains, respectively, unless a UAG stop codon was created 12 nts downstream of the Rae1 site by adding a U residue upstream of the first *gfp* codon (*spyA(UAG)-gfp*) (Fig. 4B). Similarly, the mRNA of a *bmrX-gfp* upstream fusion was insensitive to Rae1, while introduction of a stop codon 12 nts from the Rae1 site (*bmrX(UGA)-gfp*) led to a 2.9-fold Rae1-dependent destabilization, similar to that of a *gfp-bmrX* downstream fusion (3.9-fold; Fig. Sup3).

**Figure 4:**
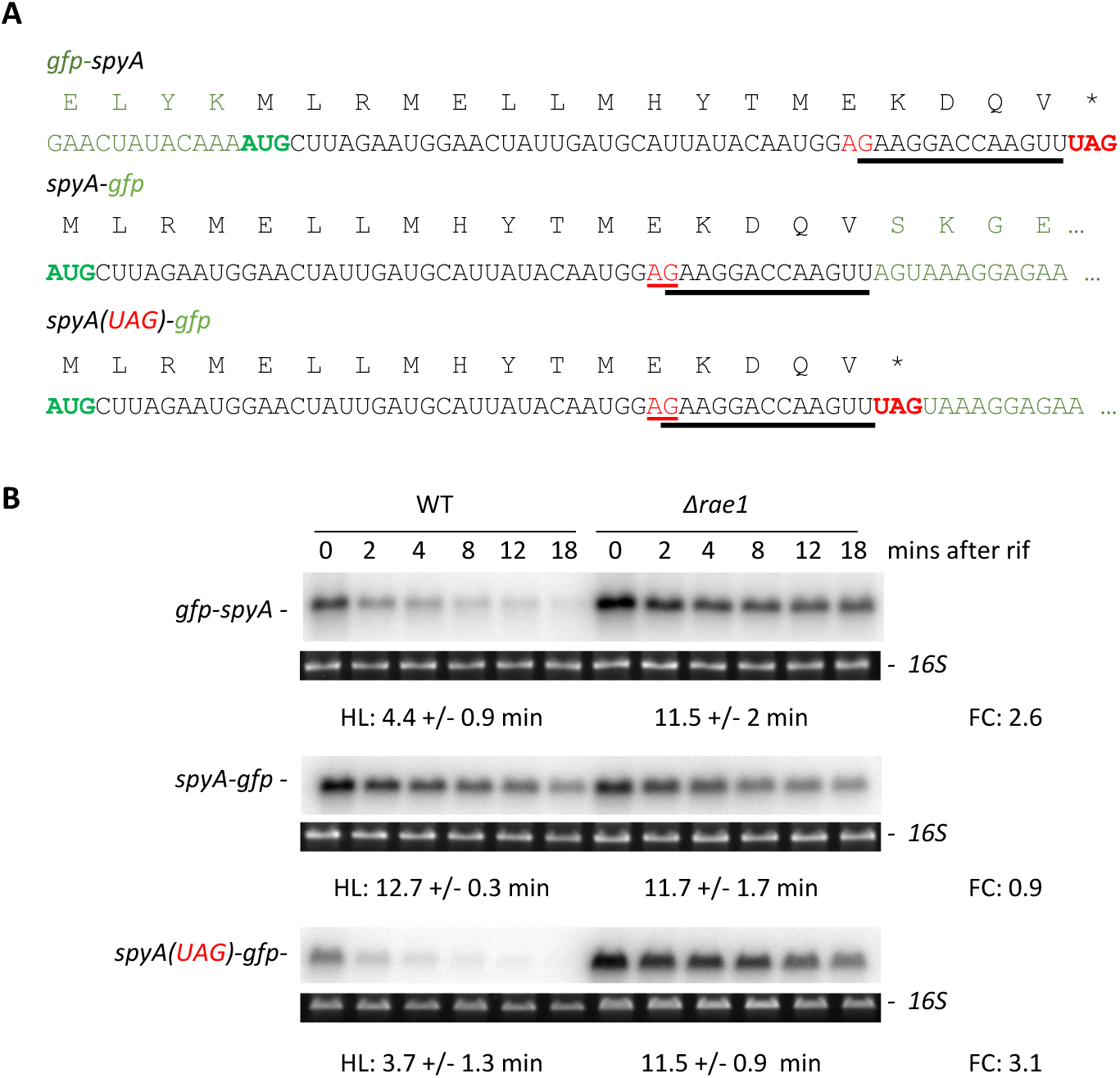
The stop codon plays a crucial role in Rae1 cleavage of *spyA*. A. Sequence of *spyA-gfp* translational fusion. The *gfp* coding sequence was fused upstream (*gfp-spyA*) or downstream (*spyA-gfp*) of the *spyA* ORF. A stop codon was placed in the *spyA(UAG)-gfp* fusion 12 nts from the Rae1 site (underlined in red). Black: *spyA* sequence. Green: *gfp s*equence. Bold green: start codon. Bold red: stop codon. B. Stability of mRNA *spyA-gfp* translational fusion revealed by Northern blot in WT and △*rae1* strains. RNAs were extracted 0, 2, 4, 8, 12, and 18 min after addition of rifampicin at 150 µg/ml. Half-lives (HL) were calculated from two biological replicates. The blots were probed with oligonucleotide CC2422 that hybridized to *gfp*. The fold-change (FC) in half-lives is given next to each autoradiogram.

Since we observed the same effect of moving the stop codon further downstream with multiple different insertion sequences in *spyA* and *bmrX* (18 nts or *gfp*), these data suggest that an appropriate distance between the Rae1 cut site and the stop codon is a key requirement for Rae1 cleavage for these two mRNAs.

### RF1 and RF2 have different impacts on Rae1 cleavage of the *spyA* mRNA

The data presented above suggested that the stop codon plays a key role in promoting efficient Rae1 cleavage of the *spyA* mRNA. We asked whether the nature of the codon itself might influence Rae1 efficiency by mutating the UAG stop codon to either UAA or UGA in the context of the *spyA* construct integrated at the *amyE* locus. The UAG codon is recognized by release factor 1 (RF1) and UGA by RF2, while the UAA codon is recognized by both release factors. The stabilities of the *spyA* stop codon mutants were measured in WT and Δ*rae1* strains by Northern blot (Fig. 5A). Although the *spyA(UAA)* and *spyA(UGA)* transcripts were still stabilized in the *rae1* strain (2.8-fold and 2.1-fold, respectively), their stabilization was weaker than in the wild-type (UAG) context (5.9-fold). Thus, the *spyA* mRNA with an RF1-specific stop codon appeared to be more sensitive to Rae1, suggesting that translation termination at this particular stop codon is least efficient in the context of the *spyA* mRNA.

**Figure 5:**
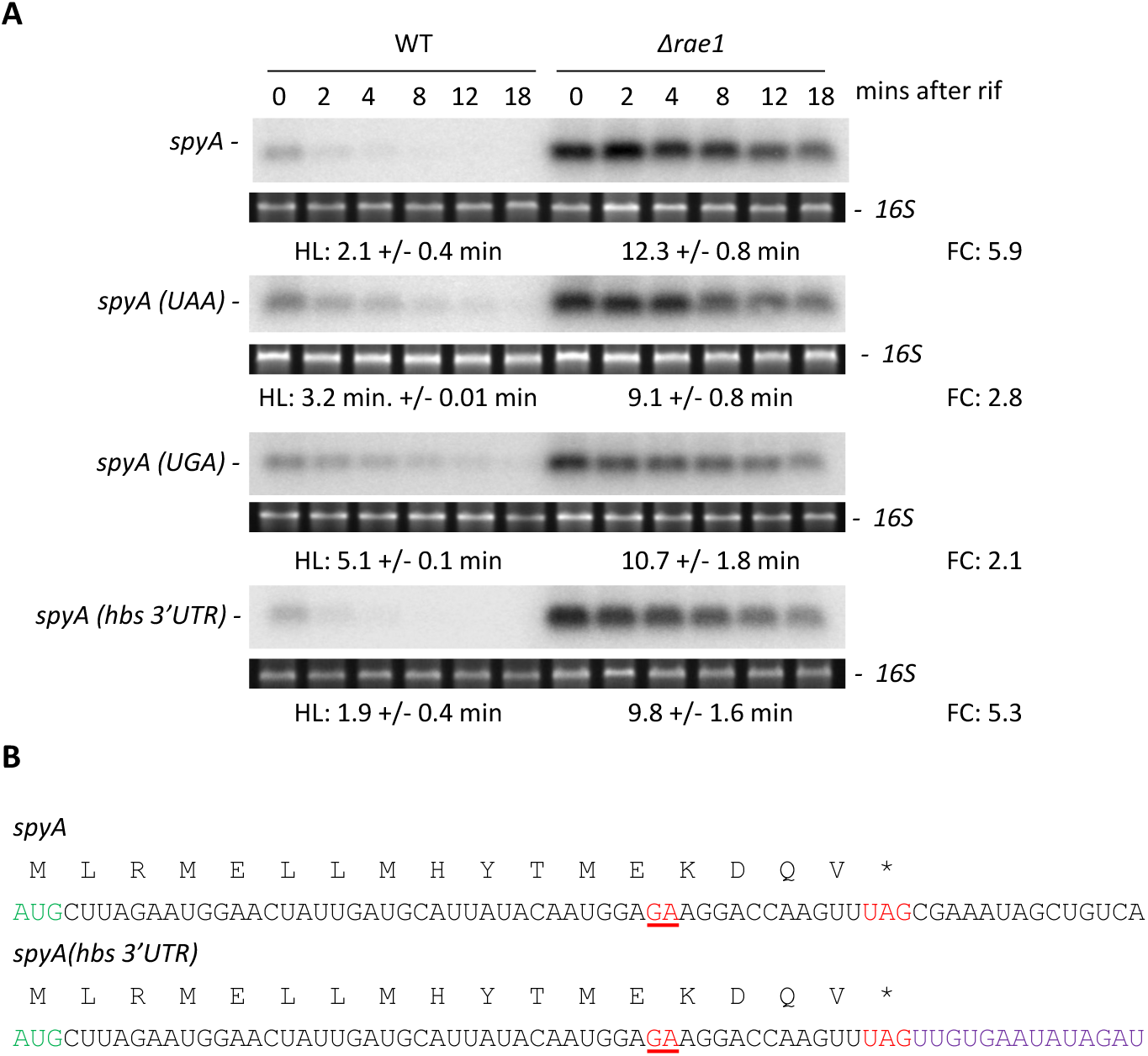
The nature of the stop codon impact Rae1 cleavage efficiency. A. Stability of the *spyA* mRNA with different stop codons or a different 3’-UTR revealed by Northern in WT and △*rae1* strains. Strains expressing the *spyA* mRNA sequence carrying either UAG, UAA or UGA stop codons, or the *hbs* 3’-UTR, were generated. RNAs were extracted 0, 2, 4, 8, 12, and 18 min after addition of rifampicin at 150 µg/ml. The blots were probed with CC2799. Half-lives (HL) were calculated from two biological replicates. The fold-change (FC) in half-lives is given next to each autoradiogram. B. In the *spyA(hbs3’UTR)* construct, the 3’UTR of *spyA* was replaced by the *hbs* 3’-UTR (in violet).

The nucleotide-context of the stop-codon, especially that immediately after the stop codon, has been shown to impact translation termination efficiency in *E. coli* ^37,38^. To investigate whether the sequence context immediately downstream of the *spyA* stop codon might impact Rae1 cleavage, we made an additional construct, *spyA* (*hbs 3’UTR*), in which the *spyA* 3’ UTR was replaced by that of the *hbs* gene (Fig. 5B). We observed that the stability of the *spyA (hbs 3’UTR)* transcript was similar to that of the WT *spyA* mRNA (1.9 vs 2.1 mins half-life in the wild-type strain, respectively), and both were similarly sensitive to Rae1 deletion (5.9 fold vs 5.3 fold, respectively), indicating that the *spyA* sequence downstream the stop codon is not required for Rae1 to promote *spyA* mRNA degradation (Fig. 5A). Consistent with this data, the *spyA, spyA UAG(+18 nts)* and *spyA(UAG)-gfp* mRNAs, which all have different sequences immediately downstream of the stop codon, are all sensitive to Rae1.

Our data above suggested that RF1 is less efficient at peptide chain release than RF2 on the *spyA* with their cognate stop codons, which might allow Rae1 more time to position itself on the ribosome to cleave the *spyA* target. To directly test the relative impacts of RF1 and RF2 on Rae1 cleavage, we translated WT *spyA* and *spyA(UGA)* transcripts in *vitro* in the presence or absence of Rae1, and *B. subtilis* RF1 or RF2 at 2 or 20 pmoles per reaction (Fig. 6). WT *spyA* and *spyA(UGA)* were both cleaved when Rae1 was added to the reaction, giving rise to the 72-nt cleavage product (Fig. 6, lanes 3 and 9). Addition of excess RF1 (20 pmol RF1 *vs.* 2.7 pmol ribosomes) to the WT *spyA* reaction did not impact Rae1 cleavage (Fig. 6, lane 5). In contrast, addition of 20 pmol RF2 inhibited the cleavage of the *spyA(UGA)* transcript (Fig. 6, lane 11). In control reactions, addition of excess RF2 did not interfere with Rae1 cleavage of the WT *spyA* transcript (Fig. 6, lane 6) and addition of excess RF1 did not interfere with Rae1 cleavage of the *spyA(UGA)* transcript (Fig. 6, lane 12), as expected. Thus, the two RFs appear to behave differently on their cognate transcripts. If we assume that addition of excess RFs should tend to increase termination efficiency and that this is inversely related to cleavage efficiency by Rae1, these results are consistent with the idea that RF1 is relatively inefficient at promoting termination at the UAG codon of the *spyA* transcript. Altogether, these data support a model in which ribosome pausing on the *spyA* mRNA, is caused by low efficiency peptide chain release by RF1 at the UAG stop codon, allowing time for Rae1 to access its target for cleavage.

**Fig 6:**
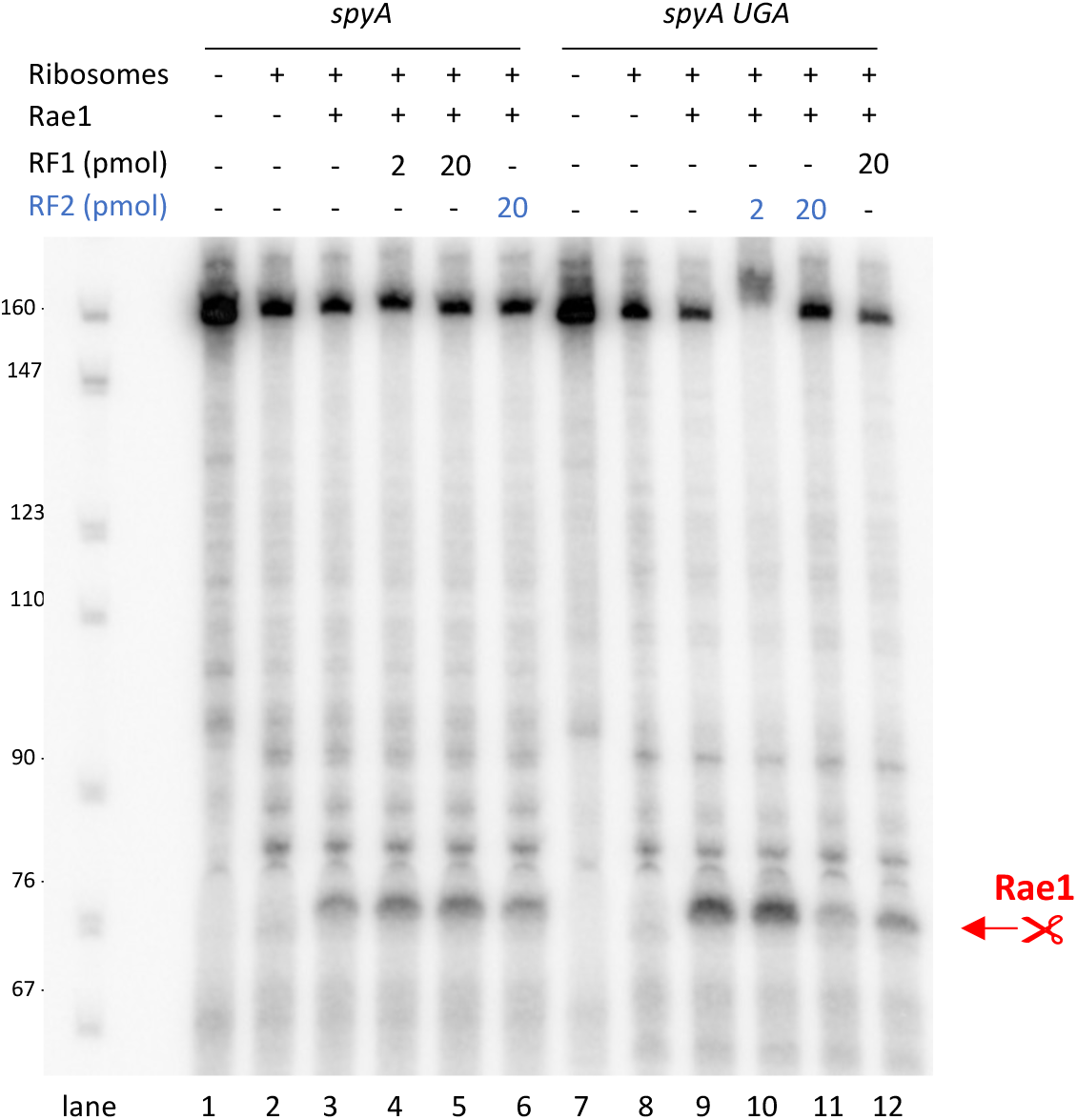
RF1 and RF2 affect Rae1 cleavage differently. *In vitro* cleavage assay of *spyA* by Rae1 in the presence of RF1 or RF2. 1 pmol of the *spyA in vitro* transcript, recognized by RF1, or *spyA(UGA)*, recognized by RF2, were incubated in presence of 2.7 pmol of ribosomes alone (lanes 1 and 7), 38 pmol of Rae1 (lanes 3 and 9) or with 2 pmol (lanes 4 and 10) or 20 pmol (lanes 5-6 and 11-12) of *B. subtilis* RF1 or RF2. The band corresponding to the transcript cleaved by Rae1 is highlighted in red.

### The SpyA peptide sequence is a key determinant of Rae1 sensitivity

We had previously shown that Rae1 cleavage is dependent on the *spyA* peptide sequence using a +1 frame-shifted (F+1) mutant retaining the nucleotide sequence encompassing the Rae1 cleavage site ^26^. However, in this construct, the Rae1 site was no longer located 12 nts upstream of a stop codon. Given what we know now about the importance of the position of the stop codon relative to the cleavage site, we could not exclude the possibility that the insensitivity of the *spyA (F+1)* transcript to Rae1 was due to the displacement of the stop codon rather than the modification of the peptide reading frame. We therefore made a new variant of the F+1 construct (*spyA-AAmod*) that restored a stop codon 12 nts downstream of the Rae1 cleavage site (Fig. 7A). Analysis of *spyA-AAmod* stability in WT and Δ*rae1*strains by Northern blot showed that this transcript was no longer sensitive to Rae1 (6.2 min and 6 mins half-lives in the WT and Δ*rae1* strains, respectively), while the WT *spyA* mRNA was stabilized 5.9-fold in the Δ*rae1* strain (Fig. 7B). We recapitulated these data by adding 5’-labelled WT *spyA* and *spyA-AAmod* mRNAs to *in vitro* translation/cleavage reactions primed with *B. subtilis* 70S ribosomes, in the presence or absence of Rae1. A Rae1 cleavage product of the expected size (72 nts) was only observed with the WT *spyA* construct, confirming that *spyA-AAmod* mRNA is not cleaved by Rae1 (Fig. 7C).

**Figure 7:**
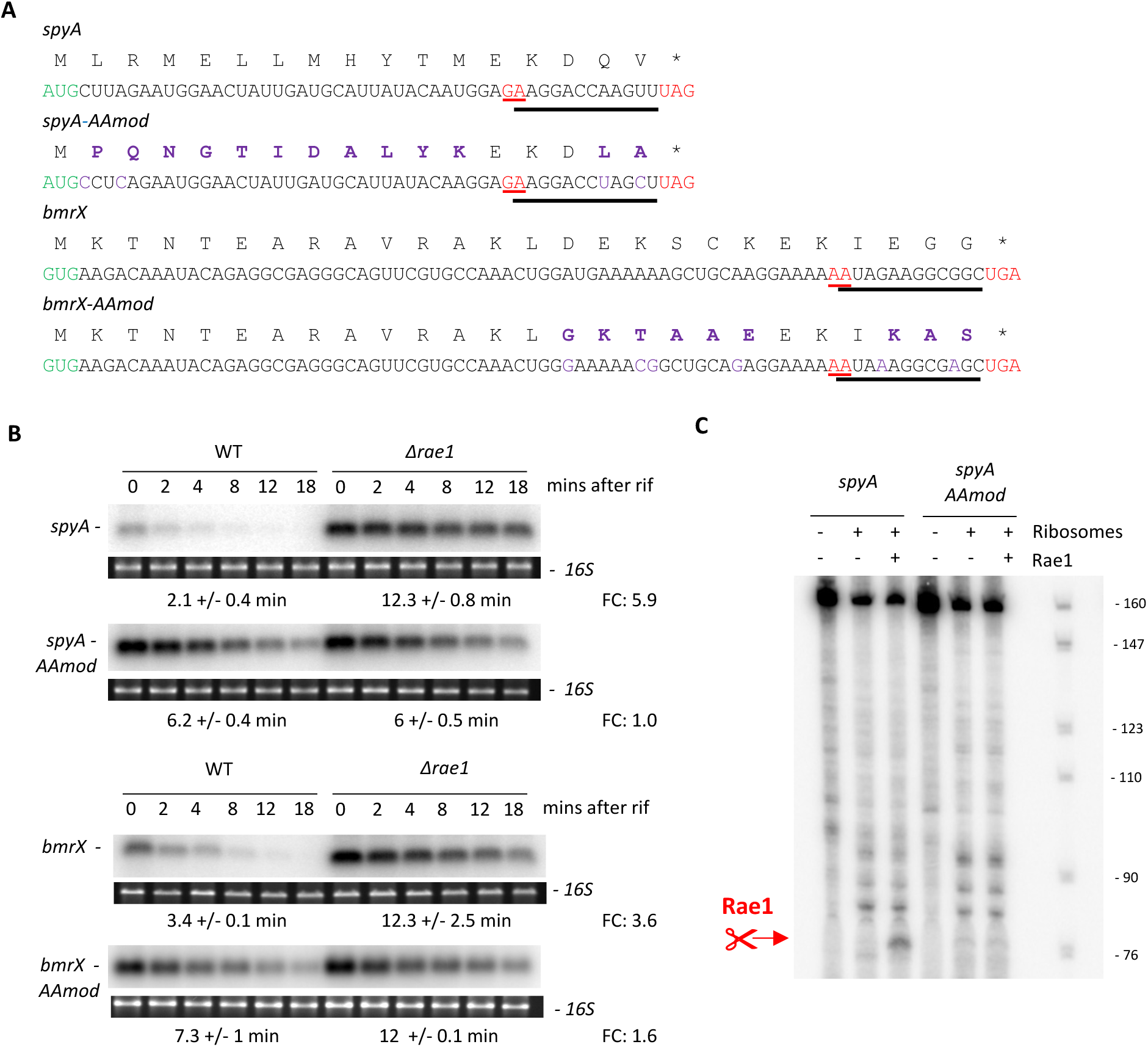
Modification of the SpyA or BmrX peptide sequence inhibits Rae1 cleavage. A. The sequences of *spyA* and *bmrX* were mutated to modify the peptide sequence without altering the 12-nt distance between the Rae1 cleavage site and the stop codon. The mutated nucleotides are shown in purple as well as the modified amino acids. Green: start codon. Red: stop codon. Red underlined: Rae1 cleavage site. B. Stability of the *spyA-AAmod* and *bmrX-AAmod* mRNA, respectively, revealed by Northern blot in the WT and △*rae1* strains. RNAs were extracted 0, 2, 4, 8, 12, and 18 min after addition of rifampicin at 150 µg/ml. The blots were probed with CC2799 or CC3271. Half-lives (HL) were calculated from two biological replicates. The fold-change (FC) in half-lives is given next to each autoradiogram. C. *In vitro* cleavage assay of *spyA* and *spyA-Aamod* mRNAs by Rae1 during translation. 1 pmol of 5’-labelled mRNAs were incubated in presence of 2.7 pmol of ribosomes and 38 pmol of Rae1. A control for RNA integrity was added without ribosomes or Rae1. The band corresponding to the transcript cleaved by Rae1 is highlighted in red.

The stability of the *bmrX-AAmod* mRNA, in which the peptide sequence was modified by a frame shift mutation, but still contained the Rae1 site 12 nts from the stop codon, was also less sensitive to Rae1 *in vivo*: 1.6-fold stabilization for *bmrX-AAmod* compared to 3.6-fold stabilization for the WT *bmr*X mRNA in the Δ*rae1* strain (Fig. 7B). These experiments highlight the influence of both the SpyA and BmrX peptide sequences in the Rae1 cleavage mechanism.

Neither the *spyA* nor *bmrX* mRNAs possess any obvious sequence features known to promote ribosome pausing, such as rare codons or polyproline motifs (Fig. 7A). However, some nascent peptide sequences can act as arrest peptides and promote ribosome stalling through interactions between specific amino acids and the peptide exit tunnel^39–41^, which can accommodate 30 to 40 amino acids ^39,42,43^. To identify the element(s) of the SpyA peptide sequence required to promote ribosome pausing and thereby Rae1 cleavage, we made a series of deletions from the beginning of the ORF, well upstream of the Rae1 cleavage site (Fig. 8A). All mutations were made in the *spyA* construct integrated at the *amyE* locus and their sensitivity to Rae1 was analyzed by measuring their stability in the WT and Δ*rae1* strains. The *spyA (16-*Δ*L)* transcript lacking the leucine codon at position 2 was still sensitive to Rae1 (4.9-fold stabilization), while the *spyA-(15-*Δ*LR)* and *spyA (14-*Δ*LRM)* transcripts lacking codons 2-3 and 2-4, respectively, were no longer Rae1-sensitive (1.3-fold and 1.1-fold stabilization, respectively) (Fig. 8B and 8C). In agreement, the *spyA (14-*Δ*LRM)* transcript was no longer cleaved *in vitro* (Fig. 8D).

**Figure 8:**
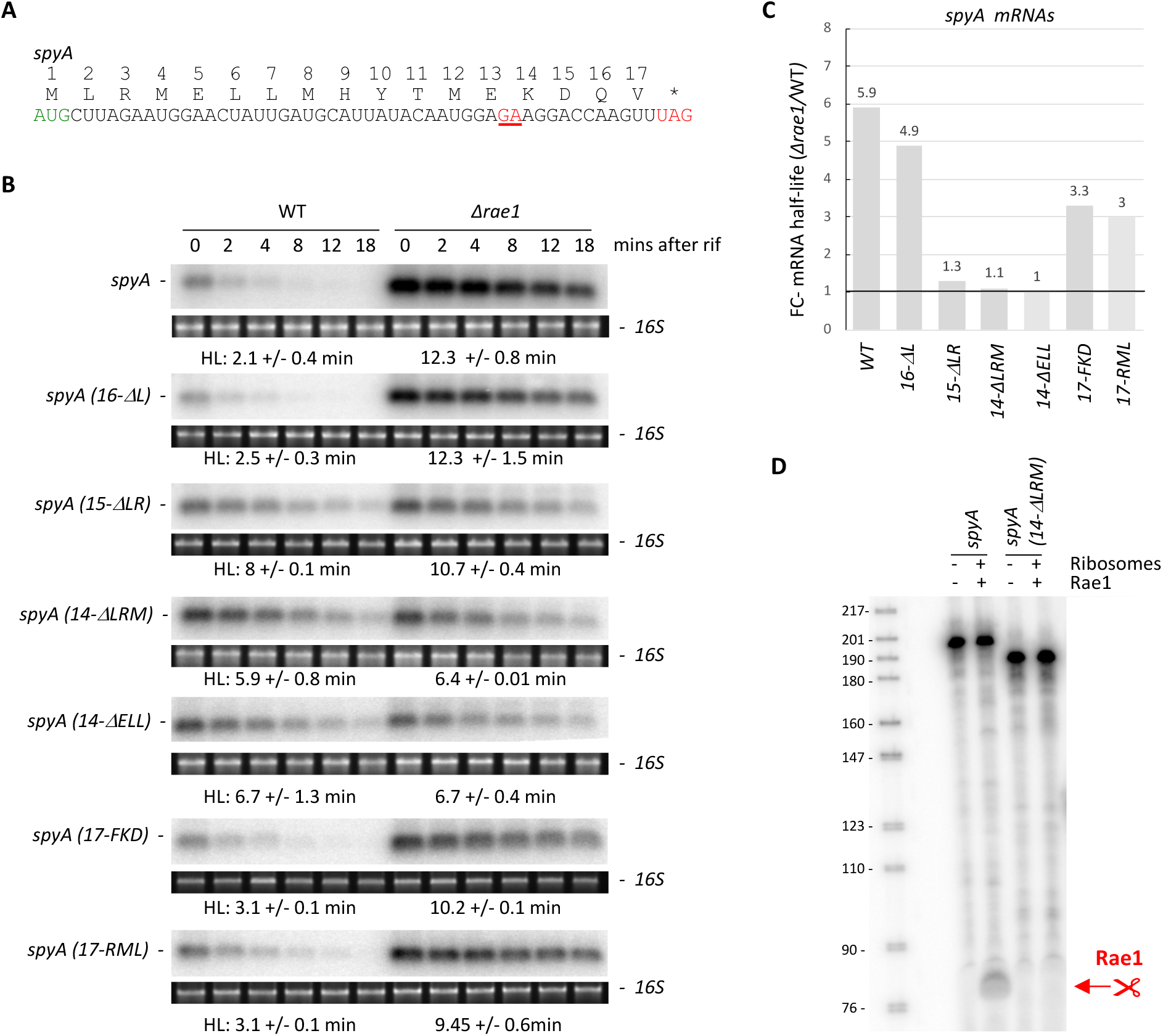
Shortening the *spyA* ORF makes the transcript insensitive to Rae1. A. Peptide and nucleotide sequence of *spyA.* The Rae1cleavage site is underlined in red, the start codon in green and the stop codon in red. B. Stability of *spyA* mRNAs of various lengths in WT and △*rae1* strains revealed by Northern Blot. RNAs were extracted 0, 2, 4, 8, 12, and 18 min after addition of rifampicin at 150 µg/ml. Half-lives (HL) were calculated using two biological replicates. The blots were probed with oligonucleotide CC2799. The *spyA* sequence (17 codons) was reduced by serial deletion from the N-terminus of 1 codon (*16-*△*L*), 2 codons (*15-*△*LR*) or 3 codons (*14-*△*LRM*) or by deletion of codons 5 to 7 (*14-*△*ELL*). The LRM amino acids from position 2 to 4 were also substituted by FKV (*17-FKV*) or RML (*17-RML*). C. Fold change (FC) of mRNA half-lives between Δ*rae1* and WT strains for the different *spyA* constructs. D. *In vitro* translation cleavage assay of *spyA* and *spyA(14-*△*LRM)* in presence of 2.7 pmol ribosomes and 38 pmol Rae1. A control for RNA integrity was added without ribosomes and Rae1. The band corresponding to the transcript cleaved by Rae1 is highlighted in red.

To determine whether specific amino acids within the ^2^LRM^4^ motif modulated the ability of Rae1 to cleave, we replaced this sequence with either FKD, *spyA (17-FKD*), or RML, *spyA (17-RML*). The *spyA (17-FKD)* and *spyA (17-RML)* mRNAs were both stabilized in the *rae1* strain (3.3-fold and 3-fold respectively), indicating that the LRM motif *per se* was not required for Rae1 cleavage, but may contribute to efficiency (Fig. 8B and 8C). To confirm these data, we deleted the codons at position 5-7, giving rise to the *spyA (14-*Δ*ELL)* transcript that still contained the LRM sequence. Like the *spyA (14-*Δ*LRM)* mRNA, the stability of the *spyA (14-*Δ*ELL)* mRNA was similar in the *rae1* and the WT strains (Fig. 8B and 8C), providing further evidence that the LRM motif is not sufficient to promote Rae1 cleavage, and suggesting the length of the ORF might be a key determinant for Rae1 cleavage. To rule out the possibility that the lack of sensitivity to Rae1 was due to an effect on translation of *spyA*, we measured the level of translation for each of these constructs by following methionine-S^35^ incorporation into peptides *in vitro,* normalized for the number of methionines in the sequence (Fig. Sup4). Although migration position of mutant peptides varied, presumably because of their different amino acid compositions, all were translated at least as efficiently as WT *spyA*. Altogether these data indicate that modification of the SpyA or BmrX peptide sequence inhibits Rae1 cleavage and suggest that shortening the *spyA* ORF to less than a critical length of 16 codons makes the transcript insensitive to Rae1.

### The ^5^ELLMHYTMEKDQV^17^ sequence is essential for Rae1-induced destabilization of *spyA*

To further investigate to what extent the length of the *spyA* ORF can affect Rae1 cleavage, we used the *gfp-spyA* translational fusion in which we replaced the wild-type *spyA* sequence by *spyA(14-*Δ*LRM),* giving rise to *gfp-spyA(14-*Δ*LRM)* (Fig. 9A) and measured its stability in WT and Δ*rae1* strains, compared to *gfp-spyA*. Both transcripts were similarly stabilized, 2.2 and 2.6-fold respectively, in the Δ*rae1* strain (Fig. 9B and 9D), consistent with the increase in ORF length. This supports our hypothesis that Rae1 cleavage requires a minimal ORF length and does not depend on the ^2^LRM^4^ sequence *per se*. Similar results were obtained with the *bmrX* mRNA. The *bmrX (16-* Δ*KTNTEARAVR)* mRNA, with only remaining 16 codons, was less sensitive to Rae1 than the WT transcript (Fig. Sup5B and 5D), while a *gfp-bmrX(11-*Δ*KTNTEARAVRAKLDE)* translational fusion containing just the last 10 amino acids of *bmrX* fused downstream of *gfp* was still stabilized in the Δ*rae1* strain (Fig. Sup5C and 5D).

**Figure 9:**
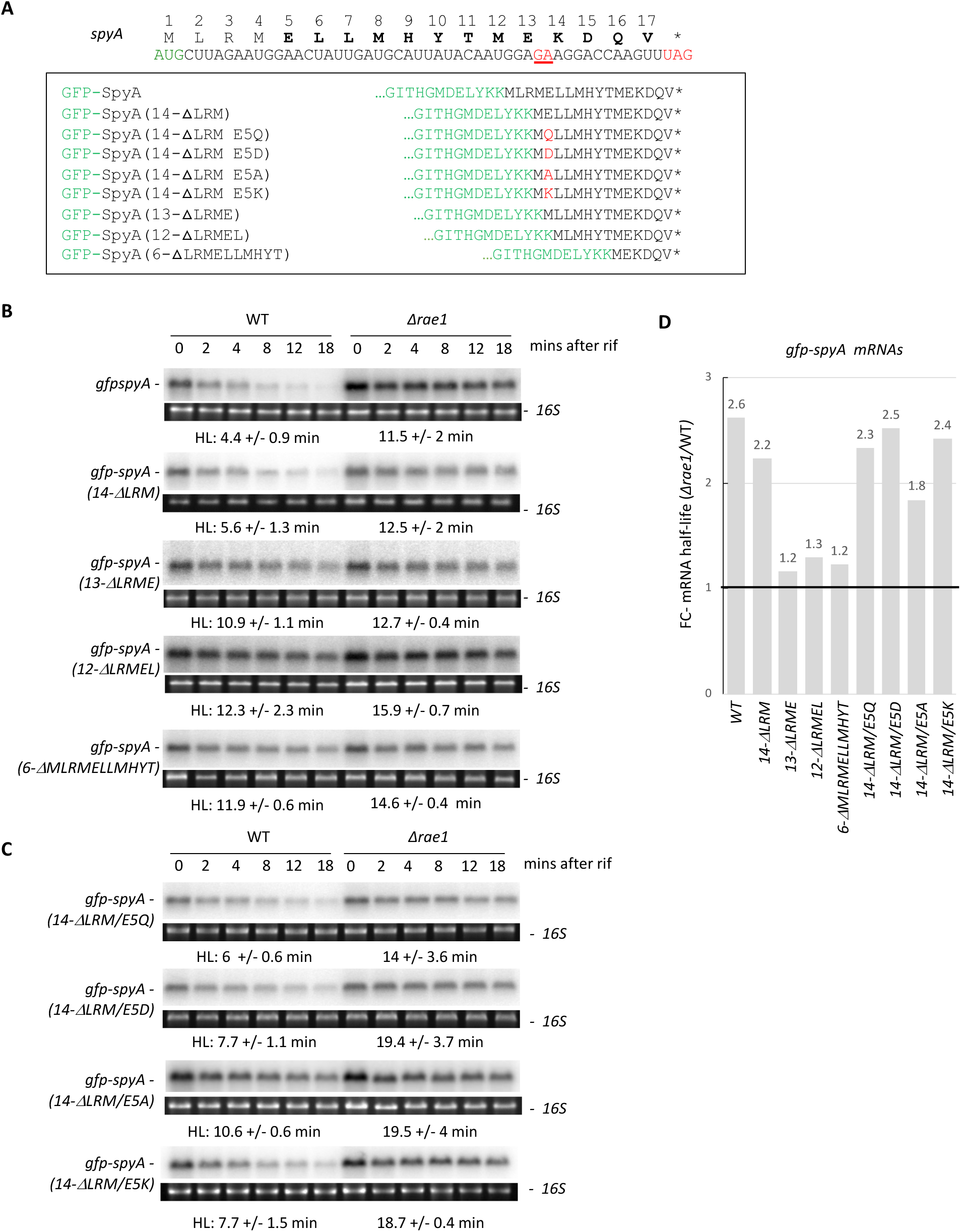
The ^5^ELLMHYTMEKDQV^17^ sequence is essential for Rae1-induced destabilization of *spyA*. A. Design of *gfp-spyA* translational fusions. Peptide and nucleotide sequence of *spyA* with the Rae1 cleavage site underlined in red, start codon in green and stop codon in red. The wild-type 17-codon sequence or shorter versions were cloned in frame downstream of the *gfp* ORF (green) and containing respectively the last 14 codons (*14-*△*LRM*), 13 codons (*13-*△*LRME*), 12 codons (*12-*△*LRMEL*) or 6 codons (*6-*△M*LRMELLMHYT*) of *spyA*. The role of glutamate at position 5 was investigated by replacing the E5 residue with an aspartate (*gfp-spyA*(*14-*△*LRM/E5D*), glutamine (*gfp-spyA*(*14-*△*LRM/E5Q*), alanine (*gfp-spyA*(*14-*△*LRM/E5A*) or lysine (*gfp-spyA*(*14-*△*LRM/E5K*). B and C. Stability of the *gfp-spyA* translational fusions with shortened *spyA* moieties in WT and △*rae1* strains revealed by Northern Blot. RNAs were extracted 0, 2, 4, 8, 12, and 18 min after addition of rifampicin at 150 µg/ml. Half-lives (HL) were calculated from biological replicates. The blots were probed with oligonucleotide CC2422. **D.** Fold change (FC) of mRNA half-lives between Δ*rae1* and WT strains for the different *spyA* constructs.

We took advantage of the *gfp-spyA* translational fusion to further investigate the role of the SpyA peptide sequence in Rae1 cleavage independently of ORF length, by deleting 4, 5 or 11 codons from the beginning of the SpyA ORF giving rise to *gfp-spyA(13-*Δ*LRME), gfp-spyA(12-*Δ*LRMEL)* and *gfp-spyA(6-*Δ*MLRMELLMHYTM)*, respectively (Fig. 9A). The three transcripts with an ORF containing the last 13, 12 and 6 codons of SpyA, respectively, all lost their sensitivity to Rae1 (Fig. 9B and 9D) suggesting that the 13-aa peptide sequence ^5^ELLMHYTMEKDQV^17^ is the minimal sequence required to promote ribosome pausing and Rae1 cleavage.

The fact that the *gfp-spyA(14-*Δ*LRM)* mRNA was sensitive to Rae1 but not the *gfp-spyA(13-*Δ*LRME)* transcript suggested that the glutamate at position 5 might be important for Rae1 cleavage. We therefore mutated the glutamate residue to aspartate (*gfp-spyA(14-*Δ*LRM/E5D)*), glutamine (*gfp-spyA(14-*Δ*LRM/E5Q)*), alanine (*gfp-spyA(14-*Δ*LRM/E5A)*) or lysine (*gfp-spyA(14-*Δ*LRM/E5K)*). Only the E5A substitution impacted the stabilization of the mRNA in the Δ*rae1* strain suggesting that it is not the charge, but rather the bulkiness of the amino acid in position 5 that contributes to ribosome stalling (Fig. 9C and 9D). Altogether these data indicate that the 13-aa peptide sequence ^5^ELLMHYTMEKDQV^17^ of SpyA and the 10-aa peptide sequence ^17^KSCKEKIEGG^26^ of BmrX are sufficient to promote mRNA degradation by Rae1.

## Discussion

The ribosome associated endoribonuclease Rae1 cleaves mRNA in a translation and reading frame dependent manner in *B. subtilis* ^26,30^. In this study, we show that Rae1 mediates mRNA cleavage in coordination with stalled ribosomes and identify multiple ORF features, in terms of both its sequence and its length, that are required for Rae1 cleavage.

The two Rae1 cleavage sites identified previously occur within the 17-aa *spyA* ORF, encoded by the polycistronic *spyTA* operon, and the cryptic 26-aa *bmrX* ORF of the polycistronic *bmrBXCD* operon ^26,30^. The cut sites were mapped between a glutamate (GAG) and a lysine codon (AAG) at positions 13 and 14 for *spyA,* and between a lysine (AAA) and an isoleucine codon (AUA) at positions 20 and 21 for *bmrX*, both located 12 nts upstream of their respective stop codons. Here, we identified a new Rae1 site within the *fliY* mRNA, located 12 nucleotides upstream of the second proline codon of a known SPP ribosome stalling motif in the *fliY* ORF. These observations converged on the notion that, at least in these three cases, Rae1 cleaves 12 nts upstream of ribosome pause sites, and were confirmed by toeprint experiments showing that ribosomes stall on the *spyA* ORF with its stop codon in the A-site and by the observation that cleavage is abolished when the stop codon is moved further downstream. We had previously proposed that Rae1 cleaves from within the A-site, based on our ability to dock the crystal structure of Rae1 there ^26^. However, these data support a new model in which Rae1 binds and cleaves at the tail end of stalled ribosomes whose A-sites are occupied by prolines or stop codons. Although not yet identified, we predict that other stalling motifs will also lead to mRNA cleavage given an appropriate sequence context for Rae1 recognition around position −12 from the A-site.

Our data suggest that ribosome stalling on the *spyA* ORF results from a low efficiency of translation termination at the stop codon, with the natural UAG stop codon promoting greater Rae1-sensitivity than a UAA or UGA stop codon. The ability of an excess of release factor RF2, but not RF1, to promote Rae1 cleavage of *spyA* constructs with their cognate stop codons *in vitro,* suggests that RF1 is inefficient at promoting termination at the *spyA* UAG termination codon. Rae1-sensitivity is clearly not restricted to the RF1/UAG combination, however, as the Rae1-dependent *bmrX* ORF has an RF2-dependent UGA stop codon. Previous examples of ribosome stalling at stop codons have been characterized from a structural point of view and may be relevant to the *spyA/bmrX* cases described here ^44–47^. For instance, tryptophan-dependent ribosome stalling at the stop codon of the 24-amino acid TnaC leader peptide in *E. coli* is promoted by free L-Trp binding and stabilization of a TnaC conformation that prevents RF2 from adopting an active conformation at the peptidyl transferase center (PTC) ^48,49^. Similarly, regulation of the expression of the Streptococcal *msrD* gene in response to macrolide exposure involves the MsrDL leader peptide causing ribosome stalling at a UAA stop codon and preventing the proper accommodation of either RF1 and RF2 ^50^.

Our results show that the amino acid sequence encoded by the *spyA* and *bmrX* ORFs is a key determinant of Rae1 cleavage. The frame-shift mutant transcript (*spyA AAmod*) that encodes a heavily modified peptide (14/17 aa), but that still contains the 9 nts surrounding the Rae1 site and the 12 nts distance between the cleavage site and the stop codon, lost its sensitivity to Rae1 both *in vivo* and *in vitro*, despite the fact that *spyA-AAmod* was translated at similar levels to WT *spyA* (Fig. Sup4). Equivalent results were obtained with the mutant *bmrX-AAmod* mRNA. We thus believe that insensitivity of the *spyA-AAmod* and *bmrX-AAmod* mRNAs to Rae1 is due to the inability of the SpyA-AAmod and BrmX-AAmod peptides to trigger ribosome pausing. Known arrest peptides, such as MifM, SecM, TnaC or MsrD, are relatively short and a ∼ 15 residue stretch appears to be sufficient to induce ribosome stalling by interacting with the exit tunnel and the PTC ^36,49–52^. Since the exit tunnel can accommodate 30 to 40 amino acids of the nascent polypeptide chain, the entire 17-aa SpyA peptide and 26-aa BmrX peptide would both be predicted to be still be inside when the ribosome reaches the stop codon in the native context ^40,42,43,53^. Studies on stalling sequences generally have highlighted the very wide variety of natural stalling motifs, making them very difficult to predict ^40,45,54^, as manifested by the absence of common motif between the SpyA and BmrX peptides. Our serial deletion approach showed that the last 13 amino acids of SpyA (^5^ELLMHYTMEKDQV^17^), and the last 10 amino acids of BmrX (^17^KSCKEKIEGG^26^), are sufficient to promote Rae1 cleavage in a context where they were fused to the C-terminus of GFP. Our data also point to a potential role for a bulky amino acid at position 5 of the SpyA peptide.

Not all stalling motifs in *B. subtilis* promote Rae1 cleavage. The *mifM* sequence, known to promote ribosome pausing to allow association of membrane transport machinery for transport of the YidC2 protein encoded downstream in *B. subtilis*, is not targeted by Rae1 (data not shown) ^36,51,55^. This suggests that at least two components are required for successful Rae1 cleavage, a peptide that provokes efficient ribosome arrest and an RNA sequence that is sensitive to Rae1 at the appropriate distance upstream of the A-site. We screened a number of other mRNAs (besides *fliY*) containing polyproline motifs for their sensitivity to Rae1 *in vivo* in the absence or EFP (Fig. S6) and did not find other obvious candidates. This could be because *B. subtilis* has at least one anti-stalling protein, YfmR, with overlapping function to EFP ^56^, or simply because a Rae1 cleavage site is not present upstream of the stall site for these mRNAs. Disentangling the exact sequence elements required for pausing versus RNA cleavage are a challenge for any given RNA and will be the target of future experiments.

Despite the overall high level of conservation of the bacterial translational machinery, there are differences between how arrest peptides interact with exit tunnel in *E. coli* vs *B. subtilis* ribosomes. The nascent chain of MifM leader peptide prevents accommodation of the incoming A-tRNA by interacting with residues of 23S rRNA and of ribosomal proteins L22 and L4 in *B. subtilis* ^36,57^. The MifM nascent chain does not promote an efficient stalling of the *E. coli* ribosome, partly due to differences between the L22 protein of these two bacteria. Interestingly, Rae1 does not cleave the *spyA* mRNA when the *in vitro* translation cleavage assay is performed with *E. coli* ribosomes (data not shown). This could be due to the inability of the SpyA peptide chain to cause *E. coli* ribosomes to stall, or the inability of Rae1 to interact with heterologous ribosomes (or both). Interactions between the nascent SpyA peptide and the component of the upper/central tunnel as well as between Rae1 and the subunits of the ribosome are under investigation by structural approaches.

An additional determinant for Rae1 cleavage is the length of the ORF. Strikingly, the last 14 codons of *spyA* were sufficient to promote Rae1-mediated destabilization in a context where the *gfp* ORF was fused upstream (*gfp-spyA(14-ΔLRM)*; but the last 15 codons of *spyA* were incapable of promoting Rae1 cleavage on their own (*spyA-(15-*Δ*LR)*. One explanation might be that at least 16 amino acids are required in the exit tunnel to trigger ribosome stalling. Another possibility is that Rae1 requires mRNAs to be covered by multiple collided ribosomes for efficient recruitment and cleavage. Based on structural studies performed with *B. subtilis* ribosomes ^28,29^, a 16-codon ORF with a ribosome stalled at the stop codon may be close to the lower limit able to accommodate two additional ribosomes in collided mode. Data from yeast studies has suggested that the architecture of three collided ribosomes provides a structural basis for preferential Hel2-mediated ubiquitinylation of trisomes compared to disomes ^58^, a property that also might be sensed by Rae1. If Rae1 does indeed detect collided ribosomes, it would appear to be independent of the known sensor of ribosome collisions, MutS2, in *B. subtilis* ^28,29^, as a *mutS2* deletion has no impact on the half-lives of either the *spyAT* or *bmrXCD* mRNAs *in vivo* ^30^.

Taken together, our data reveal a novel mechanism of post-transcriptional modulation of gene expression in *B. subtilis* by Rae1, which exploits ribosome stalling to cleave mRNAs. In the case of the *spy* and *bmr* operons, this mechanism has clearly been capitalized upon to shut down expression of the SpyT toxin ^32^ and readthrough of the *bmrB* attenuator, which would lead to futile expression of the BmrCD efflux pump in the absence of antibiotics ^30^. However, our observation that the *fliY* mRNA is also a substrate leads us to suspect that Rae1 may play a broader role in quality control elimination of other mRNAs that are encountering translation problems. Such mRNAs will be cleaved if a Rae1-sensitive site is located at the tail end of a stalled ribosome, and one of our future goals is to determine the breadth of the sequence requirements for Rae1 recognition. The wide conservation of Rae1 among Firmicutes, Cyanobacteria and the higher plant chloroplasts ^31^, far beyond the conservation of the *spy* and *bmrX* operons, further supports the idea that this enzyme may play a general role in translational quality control, probably in response to different stress conditions that remain to be defined.

## Acknowledgments

We thank lab members for helpful discussions. We are grateful to Dr. Joel Belasco for helpful discussions and for comments on the manuscript. We also thank Dr. Shinobu Chiba for the gift of the two plasmids allowing the expression of the RF1-His6 and RF2-His6 hexahistidine-tagged proteins from *B. subtilis*.

## Author contribution

VD, AD and LG performed the experiments. CC helped supervise the project and wrote the manuscript. FB contributed to the execution of the experiments, supervised the project and wrote the manuscript.

