## supplemental figures for "The endoribonuclease Rae1 from *Bacillus subtilis* cleaves mRNA upstream of stalled ribosomes"

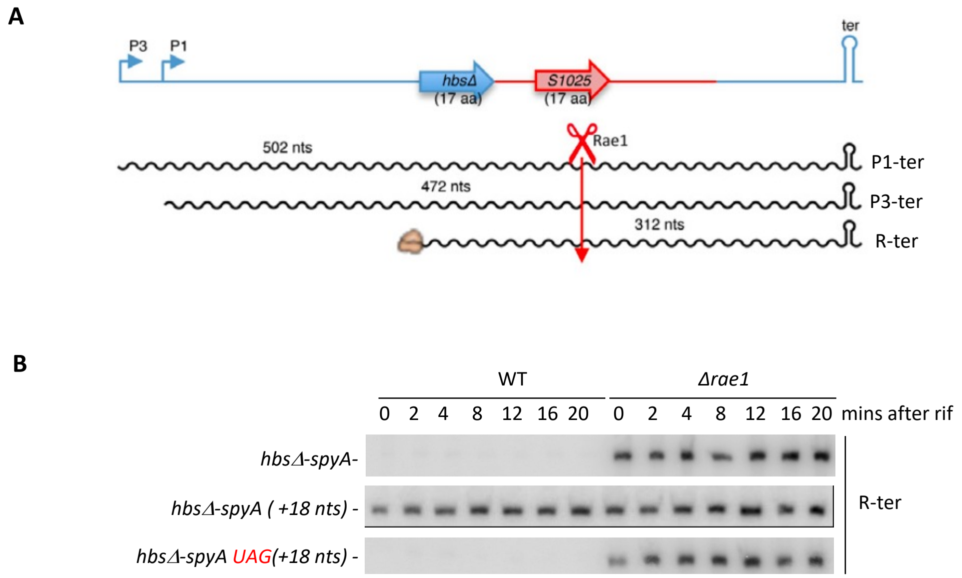


**Fig. Sup1: The distance between the Rae1 site and the stop codon is key for *spyA* mRNA cleavage by Rae1.**

**A.** Schematic of various *hbsΔ-spyA* transcripts. The *hbsΔ-spyA* construct expressed from the *amyE* locus, yields three transcripts corresponding to expression from two promoters P3 (P3-ter) and P1 (P1-ter) and a highly stable ribosome-protected species (R-ter). The *spyA* sequence is shown in red and Rae1 depicted with a scissors symbol (Leroy et al, 2017). **B.** Stability of the *hbsΔ-spyA* , *hbsΔ-spyA (+18 nts), hbsΔ-spyA UAG(+18 nts)* determined by Northern Blot. The 18-nucleotide random sequence was added downstream of the *spyA* ORF (*hbsΔ-spyA (+18 nts))* and a stop codon was recreated at its native location in this context (*hbsΔ-spyA UAG(+18 nts))*. Total RNA extracted 0, 2, 4, 8, 12, 16 and 20 min after addition of rifampicin at 150 µg/ml, in WT and the *Δrae1* strains. For simplicity, only the band corresponding to R-ter is shown. The blots were probed with CC2799 that hybridizes to *spyA*.

**A**

*bmrX*

```
 M  K  T  N  T  E  A  R  A  V  R  A  K  L  D  E  K  S  C  K  E  K  I  E  G  G  *
GUGAAGACAAAUACAGAGGCGAGGGCAGUUCGUGCCAAACUGGAUGAAAAAAGCUGCAAGGAAAAAAUAGAAGGCGGCUGA
```

12 nts

*bmrX (+18 nts)*

```
 M  K  T  N  T  E  A  R  A  V  R  A  K  L  D  E  K  S  C  K  E  K  I  E  G  G  S  K  G  E  E  L *
GUGAAGACAAAUACAGAGGCGAGGGCAGUUCGUGCCAAACUGGAUGAAAAAAGCUGCAAGGAAAAAAUAGAAGGCGGCAGUAAAGGAGAAGAACUUUGA
```

18 nts

*bmrXUGA(+18 nts)*

```
 M  K  T  N  T  E  A  R  A  V  R  A  K  L  D  E  K  S  C  K  E  K  I  E  G  G  *
GUGAAGACAAAUACAGAGGCGAGGGCAGUUCGUGCCAAACUGGAUGAAAAAAGCUGCAAGGAAAAAAUAGAAGGCGGCUGAGUAAAGGAGAAGAACUUUGA
```

12 nts

**B**

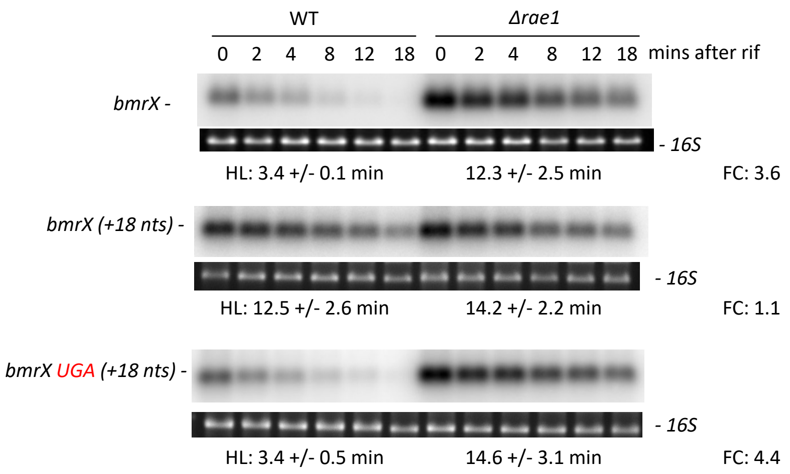


**Fig. Sup2 : The distance between the Rae1 cleavage site and the stop codon is important for *bmrX* mRNA cleavage by Rae1**

**A.** Localization of Rae1 cleavage site on *bmrX* mRNAs. The Rae1 cleavage site is underlined in red. The nucleotide distance between the stop codon and the Rae1 site is represented by a black line. Start codons are highlighted in green, stop codons in red and the 18-nt random sequence added downstream of *bmrX* mRNA in blue (*bmrX (+18 nts))*. In the *bmrX UGA(+18 nts)* construct, a stop codon was reintroduced at its native location. **B.** Stability of various *bmrX* transcripts in WT and *Δrae1* strains revealed by Northern Blot. Total RNA was extracted 0, 2, 4, 8, 12 and 18 min after addition of rifampicin at 150 µg/ml and the blots were probed with oligonucleotide CC3271. The half-lives (HL) of *bmrX (+18 nts)* and *bmrX UGA(+18 nts*) were calculated from two biological replicates. The fold-change (FC) in half-lives is given next to each autoradiogram.

**A**

*gfp-bmrX*

E L Y K V K T N T E A R A V R A K L D E K S C K E K I E G G *

GAACUAUACAAA**GUG**AAGACAAAUACAGAGGCGAGGGCAGUUCGUGCCAAACUGGAUGAAAAAAGCUGCAAGGAAAAAAUAGAAGGCGGC**UGA**

*bmrX-gfp*

M K T N T E A R A V R A K L D E K S C K E K I E G G S K G E …

**GUG**AAGACAAAUACAGAGGCGAGGGCAGUUCGUGCCAAACUGGAUGAAAAAAGCUGCAAGGAAAAAAUAGAAGGCGGCAGUAAAGGAGAA …

*bmrX(UGA)-gfp*

M K T N T E A R A V R A K L D E K S C K E K I E G G *

**GUG**AAGACAAAUACAGAGGCGAGGGCAGUUCGUGCCAAACUGGAUGAAAAAAGCUGCAAGGAAAAAAUAGAAGGCGGC**UGA**GUAAAGGAGAA …

**B**

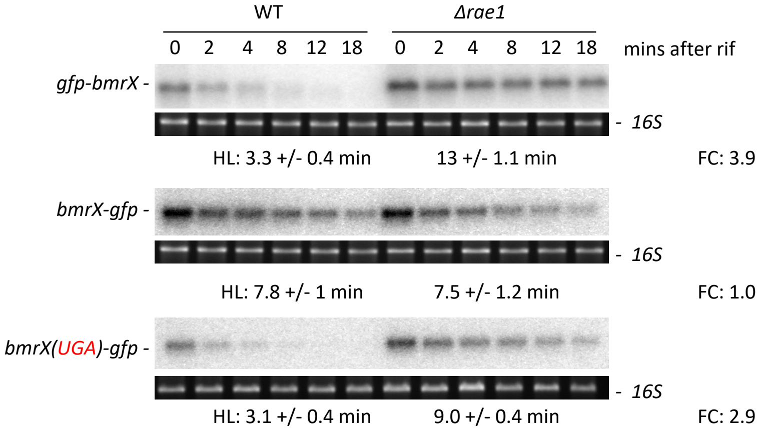


**Fig. Sup3: Rae1 cleavage of the *bmrX* mRNA requires a stop codon**

**A.** Sequence of *bmrX-gfp* translational fusion. The GFP coding sequence was fused upstream (*gfp-bmrX*) or downstream (*bmrX-gfp*) of the *bmrA* ORF. In the *bmrX(UGA)-gfp* fusion, a stop codon was placed 12 nts from the Rae1 site (underlined in red). Black: *bmrX* sequence. Green: *gfp s*equence. Bold green: start codon. Bold red: stop codon. **B.** Stability of *gfp-bmrX, bmrX-gfp* and *bmrX(UGA)-gfp* transcripts revealed by Northern blot in WT and *Δrae1* strains. RNAs were extracted 0, 2, 4, 8, 12, and 18 min after addition of rifampicin at 150 µg/ml. The blots were probed with CC2422 that hybridized to *gfp*. The calculated half-lives (HL) are the average of at least two independent experiments. The fold-change (FC) in half-lives is given next to each autoradiogram.

.

**A**

SpyA WT
**M** L R **M** E L L **M** H Y T **M** E K D Q V
SpyA AAmod
**M** P Q N G T I D A L Y K E K D L A
SpyA (15-ΔL)
**M** **M** E L L **M** H Y T **M** E K D Q V
SpyA (16-ΔL)
**M** R **M** E L L **M** H Y T **M** E K D Q V
SpyA(14-ΔLRM)
**M** E L L **M** H Y T **M** E K D Q V
SpyA(14-ΔELL)
**M** L R **M** E H Y T **M** E K D Q V
SpyA(17-FKD)
**M** F K D E L L **M** H Y T **M** E K D Q V
SpyA (17-RML)
**M** R **M** L E L L **M** H Y T **M** E K D Q V

**B**

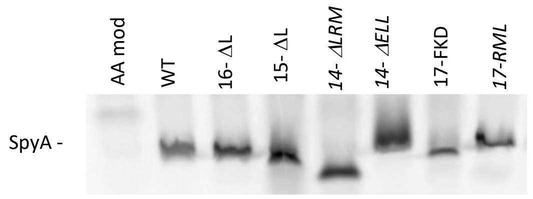


**C**

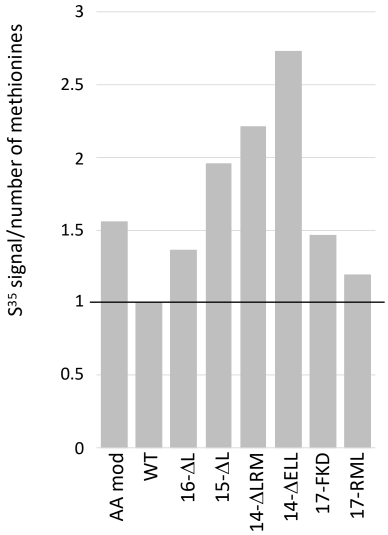


**Fig. Sup4**. **Measurement of SpyA peptide synthesis**

**A.** Sequence of the various SpyA peptides, with methionines highlighted in bold. The FKD and RML mutations are underlined. **B.** $S^{35}$-methionine-labelled *in vitro* translation reactions were performed in PURExpress cell-free extracts with 2.7 pmol *B. subtilis* ribosomes and a PCR template to transcribe *spyA* mRNA by T7 RNA polymerase. Peptides were run on 20% SDS-PAGE and quantified by PhosphorImager. **C.** Quantification of methionine-$S^{35}$ incorporated into SpyA peptides, normalized to the number of encoded methionines.

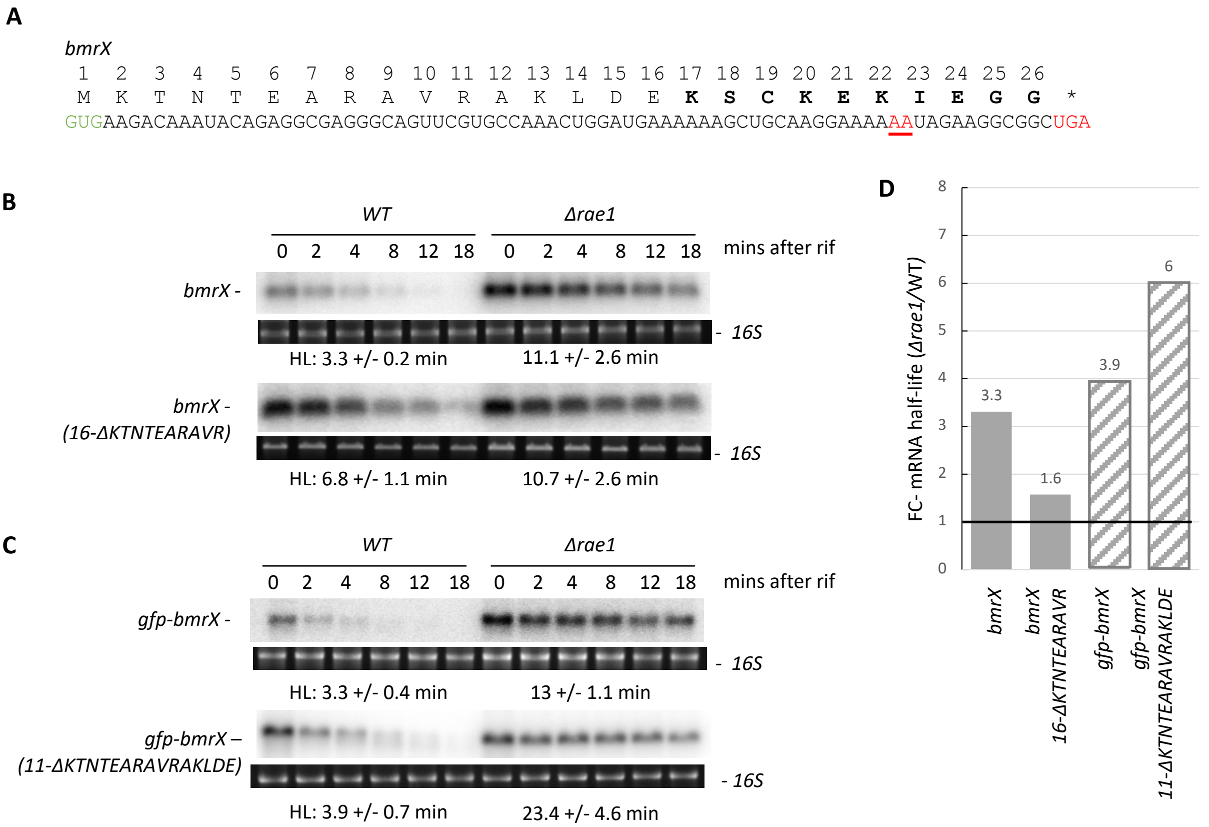


**Fig. Sup5: The [17]KSCKEKIEGG[26] sequence is sufficient for Rae1-induced destabilization of *bmrX***

**A.** Peptide and nucleotide sequences of *bmrX* with the Rae1cleavage site underlined in red, start codon in green and stop codon in red. **B.** Stability of the wild-type *bmrX* and the shorter *bmrX(16-ΔKTNTEARAVR)* mRNAs in a WT and *Δrae1* strains revealed by Northern Blot. RNAs were extracted 0, 2, 4, 8, 12, and 18 min after addition of rifampicin at 150 µg/ml. The blots were probed with oligonucleotides CC2194 or CC3271. The calculated half-lives (HL) are the average of at least two independent experiments. **C.** Stability of the *gfp-bmrX* and *gfp-bmrX(11-ΔKTNTEARAVRAKLDE)* translational fusions in WT and *Δrae1* strains revealed by Northern Blot. RNAs were extracted 0, 2, 4, 8, 12, and 18 min after addition of rifampicin at 150 µg/ml. The blots were probed with CC2422. The calculated half-lives (HL) are the average of at least two independent experiments. **D.** Fold change (FC) of mRNA half-lives between *rae1* and WT strains for the different *bmrX* constructs.

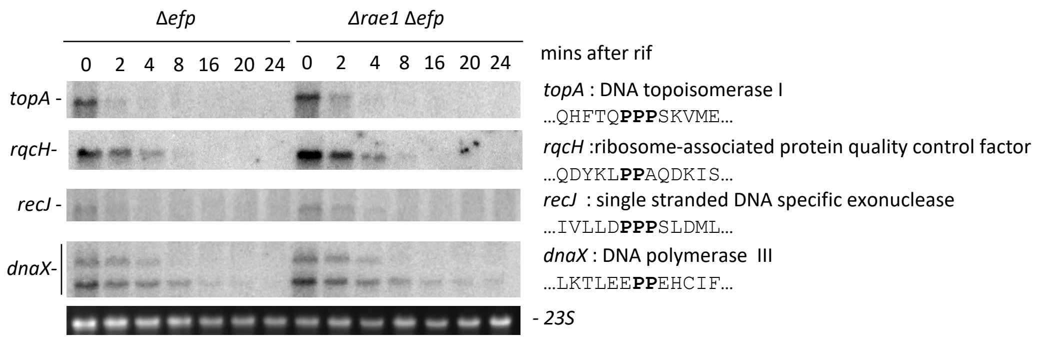


**Fig Sup6: A polyroline motif is not sufficient to trigger mRNA degradation by Rae1**

Stability of *topA*, *rqcH*, *recJ* and *dnaX* transcripts in *Δefp* and *ΔefpΔrae1* strains revealed by Northern Blot. The selected mRNAs contain different polyproline motifs within their coding sequence. RNAs were extracted after 0, 2, 4, 8, 16, 20 and 24 min after addition of rifampicin at 150µg/ml. The probes used are listed in table S1.
