## supplemental table for "The endoribonuclease Rae1 from *Bacillus subtilis* cleaves mRNA upstream of stalled ribosomes"

**Table S1.** Oligonucleotides used in this study.

| Oligo | Gene | Sequence |
| --- | --- | --- |
| CC429 | *hbs* | GTAAGCTAGATGCATACACGATCTATATTCAC |
| CC572 | *hbs* | AAGCAAAGGGATCCCCTGTCAGGAAGCCCATGCTC |
| CC1607 | *hbs* | GGTGAATTCCCGAGCTGTAATACCGGTAGACCTC |
| CC1660 | *spyA* | CAGTAATACGACTCACTATAGAAGCAACGAAAAAAACTGAAGGGG |
| CC1661 | *spyA* | GCCATCACCTGCCTCACAATAAC |
| CC1787 | pHM2 | CTTATCTTGATAATAAGGGTAAC |
| CC1808 | *spyA* | ATAAAGCTTAAACTGAAGGGGGAATGGCATATGC |
| CC2344 | *bmrC* | CGCAATCGTATAGCGCAGCCAGTACGCC |
| CC2422 | *gfp* | caccttcaaacttgacttcagcacgtgtc |
| CC2634 | *gfp* | ATTAAGCTTGAAGGGGGAATGGCATAtgagtaaaggagaagaacttttcactggag |
| CC2696 | *fliY* | ATTAAGCTTAGATCACGACATAAGAGGTGAACAAGATGG |
| CC2697 | *fliY* | AATGGATCCAAAAAGGCCATCCGTCAGGATGGCCATAATCTATCTCTCTCCTTGTGTTTCG |
| CC2799 | *spyA* | AACTTGGTCCTTCTCCATTGTATAATGC |
| CC2800 | *spyA* | CAATGGAGAAGGACCAAGTTAGTAAAGGAGAAGAACTTTTCACTGG |
| CC2813 | *spyA* | TGCGACTACTGCAGCGACAACTTGGTCCTTCTCCATTGTATAATGCATC |
| CC2814 | *spyA* | GTCGCTGCAGTAGTCGCATAGCGAAATAGCTGTCAAGAAAATGCGCGCC |
| CC2840 | *fliY* | CACTTCAATCGCTACATACGGTTCAGGAAACGC |
| CC2888 | *bmrX* | ATAAAGCTTAAACTGAAGGGGGAATGGCATGTGAAGACAAATACAGAGGCGAGGGC |
| CC2905 | *topA* | CAACGGATTGCACGCGTCCTGCGCTAAGCCC |
| CC2907 | *recJ* | CGTTCGCTTCTCCGATGTCTCCGCCTGACAGC |
| CC2910 | *bmrX* | CCCTCGCCTCTGTATTTGTCTTCACTTTGTATAGTTCATCCATGCCATGTGTAATCCC |
| CC2911 | *bmrX* | GTGAAGACAAATACAGAGGCGAGGGCAGTTCGTG |
| CC2922 | *fliY* | CAGGAATACGGTCTTACCCCAGTAAAAAAATTGATATTTCAGCTGCTCGGGTTGAATTGCTGGATG |
| CC2923 | *fliY* | CATCCAGCAATTCAACCCGAGCAGCTGAAATATCAATTTTTTTACTG |
| CC2947 | *fliY* | CATCATCAGGAATACGGTCTGTACCC |
| CC2950 | *fliY* | AATGGATCCAAAAAGGCCATCCGTCAGGATGGCCAGCATTTCATCAT |
| CC2953 | *spyA* | GCGACTACTGCAGCGACTAAACTTGGTCCTTCTCCATTG |
| CC2954 | *spyA* | CAATGGAGAAGGACCAAGTTTAGTCGCTGCAGTAGTCGC |
| CC2960 | *dnaX* | GTTTGCCCTCCTCCGTTATTCATCATTCCC |
| CC2962 | *spyA* | GGATCCGAAAAGCCCCTTAGAAAGGGGCTTTTCATTTCAATGACATTAGACTGCTAATAAAAGGAC |
| CC2993 | *spyA* | ATGCATCAATAGTTCCATATGCCATTCCCCCTTCAG |
| CC2994 | *spyA* | ATGGAACTATTGATGCATTATACAATGGAG |
| CC2997 | *spyA* | AACTTGGTCCTTCTCCATATGCCATTCCCCCTTCAG |
| CC2998 | *spyA* | ATGGAGAAGGACCAAGTTTAGCGAAATAGC |
| CC2999 | *spyA* | CTAAACTTGGTCCTTCTCCATTG |
| CC3000 | *spyA* | GGAGAAGGACCAAGTTTAGTTGTGAATATAGATCGTGTATGCATCTAGC |
| CC3001 | *spyA* | CTTGACAGCTATTTCGTCAAACTTGGTCCTTCTCC |
| CC3002 | *spyA* | GGAGAAGGACCAAGTTTGACGAAATAGCTGTCAAG |
| CC3029 | *spyA* | ATGCATCAATAGTTCATCCTTGAACATATGCCATTCCCCCTTCAG |
| CC3030 | *spyA* | GAACTATTGATGCATTATACAATGGAGAAGGACC |
| CC3031 | *spyA* | ATGCATCAATAGTTCAAGCATTCTCATATGCCATTCCCCCTTCAG |
| CC3033 | *spyA* | ATGCATCAATAGTTCCATTCTCATATGCCATTCCCCCTTCAG |
| CC3034 | *spyA* | ATGCATCAATAGTTCCATCATATGCCATTCCCCCTTCAG |
| CC3035 | *spyA* | CTCCATTGTATAATGTTCCATTCTAAGCATATGCCATTCCCCCTTCAG |
| CC3036 | *spyA* | CATTATACAATGGAGAAGGACCAAGTTTAGCG |
| CC3051 | *rqcH* | CTTGGCCGGGCAGTACTGTACGGTAGCTGTTC |
| CC3182 | *spyA* | CTTGACAGCTATTTCGTTAAACTTGGTCCTTCTCC |
| CC3183 | *spyA* | GGAGAAGGACCAAGTTTAACGAAATAGCTGTCAAG |
| CC3184 | *spyA* | CAATGGAGAAGGACCAAGTTTAGTAAAGGAGAAGAACTTTTCACTGG |
| CC3219 | *gfp* | CATTTTGTATAGTTCATCCATGCC |
| CC3220 | *spyA* | ggcatggatgaactatacaaaATGGAACTATTGATGCATTATACAATGG |
| CC3271 | *spyA* | CATATGCCATTCCCCCTTCAGTTTTTTTCGTTGC |
| CC3273 | *spyA* | GGCATGGATGAACTATACAAAATGGAGAAGGACCAAGTTTAGCGAAATAGCTGTCAAG |
| CC3274 | *bmrX* | GTGAAAAGTTCTTCTCCTTTACTCAGCCGCCTTCTATTTTTTCC |
| CC3275 | *bmrX* | GGAAAAAATAGAAGGCGGCTGAGTAAAGGAGAAGAACTTTTCAC |
| CC3305 | *spyA* | CAAGGAGAAGGACCTAGCTTAGCGAAATAGCTGTCAAG |
| CC3306 | *spyA* | CTAAGCTAGGTCCTTCTCCTTGTATAATGCATCAATAGTTCC |
| CC3314 | *spyA* | GGCATGGATGAACTATACAAAATGTTGATGCATTATACAATGGAGAAGGAC |
| CC3329 | *spyA* | GGCATGGATGAACTATACAAAATGCTATTGATGCATTATACAATGGAGAAGGAC |
| CC3332 | *bmrX* | ATGCCATTCCCCCTTCAGTTTTTTTCGTTGC |
| CC3333 | *bmrX* | GCAACGAAAAAAACTGAAGGGGGAATGGCATGTGGCCAAACTGGATGAAAAAAGCTGCAAGG |
| CC3366 | *spyA* | ggcatggatgaactatacaaaATGCAACTATTGATGCATTATACAATGG |
| CC3367 | *spyA* | ggcatggatgaactatacaaaATGGACCTATTGATGCATTATACAATGG |
| CC3481 | *bmrX* | CGTCAGCCGCAGTCTATTTTTCCTTGCAGCTTTTTTACATCC |
| CC3482 | *bmrX* | GGATGTAAAAAAGCTGCAAGGAAAAATAGACTGCGGCTGACG |
| CC3483 | *bmrX* | TCACCTCGCCTCTGTATTTGTGCCGCCTTCTATTTTTTCCTTGCAGC |
| CC3484 | *bmrX* | ACAAATACAGAGGCGAGGTGACGAAATAGCTGTCAAGAAAATGCGCG |
| CC3485 | *bmrX* | TCACCTCGCCTCTGTATTTGTTCAGCCGCCTTCTATTTTTTCCTTGCAGC |
| CC3486 | *bmrX* | TGAACAAATACAGAGGCGAGGTGACGAAATAGCTGTCAAGAAAATGCGCGC |
| CC3487 | *bmrX* | CTTGACAGCTATTTCGCTAGCCGCCTTCTATTTTTTCCTTGC |
| CC3488 | *bmrX* | GCAAGGAAAAAATAGAAGGCGGCTAGCGAAATAGCTGTCAAG |
| CC3512 | *spyA* | GGCATGGATGAACTATACAAAATGGCACTATTGATGCATTATACAATGGAG |
| CC3513 | *spyA* | GGCATGGATGAACTATACAAAATGAAACTATTGATGCATTATACAATGG |
| CC3534 | *bmrX* | GGCATGGATGAACTATACAAAGTGAAAAGCTGCAAGGAAAAAATAGAAGGCGGCT |
| HP10 | pHM2 | CTCTTCGCTATTACGCCAGC |

**Table S2.** Bacterial strains used in this study

| **Strain** | **Genotype** | **Reference** | | |
| --- | --- | --- | --- | --- |
| ***E. coli*** |  | |  | |
| CCE223 | BL21 CodonPlus pRIL + pET28-Rae1CHis | | | ^26^ |
| CCE300 | BL21 CodonPlus pRIL + pET28-RF1His_6_ | | | This study |
| CCE299 | BL21 CodonPlus pRIL + pET28-RF2His_6_ | | | This study |
| ***B. subtilis*** |  | | |  |
| SSB1002 | W168 *trpC+* | | | Lab strain |
| CCB375 | W168 *rae1::pMUTIN ery* | | | ^26^ |
| CCB434 | W168 *rnjA::spc* | | | ^65^ |
| CCB841 | W168 *amyE::pHM2-hbsΔ-spyA* (pl707) | | | ^26^ |
| CCB843 | W168 *rae1::pMUTIN ery amyE::pHM2-hbsΔ-spyA* (pl707) | | | ^26^ |
| CCB1350 | W168 *amyE::pHM2-gfpspyA* (pl869) | | | This study |
| CCB1351 | W168 *rae1::pMUTIN ery amyE::pHM2-gfpspyA* (pl869) | | | This study |
| CCB1450 | W168 *amyE::pHM2-hbsΔ-spyA+18nts* (pl895) | | | This study |
| CCB1456 | W168 *pHM2-fliY* (pl856) | | | This study |
| CCB1461 | W168 *rae1::pMUTIN ery amyE::pHM2-fliY* (pl856) | | | This study |
| CCB1463 | W168 *efp::kan* (BKK24450) | | | ^59^ |
| CCB1470 | W168 *amyE::pHM2-spyAgfp* (pl899) | | | This study |
| CCB1471 | W168 *rae1::pMUTIN ery amyE::pHM2-spyAgfp* (pl899) | | | This study |
| CCB1476 | W168 *amyE::pHM2-bmrXgfp* (pl904) | | | This study |
| CCB1477 | W168 *rae1::pMUTIN ery* *amyE::pHM2-bmrXgfp* (pl904) | | | This study |
| CCB1479 | W168 *efp::kan amyE::pHM2-fliY* (pl856) | | | This study |
| CCB1480 | W168 *rae1::pMUTIN ery efp::kan amyE::pHM2-fliY* (pl856) | | | This study |
| CCB1481 | W168 *efp::kan* | | | This study |
| CCB1482 | W168 *rae1::pMUTIN ery efp::kan* | | | This study |
| CCB1550 | W168 *efp::kan* *amyE::pHM2-ΔfliYPP* (pl922) | | | This study |
| CCB1551 | W168 *rae1::pMUTIN ery efp::kan amyE::pHM2-ΔfliYPP* (pl922) | | | This study |
| CCB1559 | W168 *efp::kan*  *amyE::pHM2-ΔfliYAA* (pl923) | | | This study |
| CCB1564 | W168 *rae1::pMUTIN ery efp::kan amyE::pHM2-ΔfliYAA* (pl923) | | | This study |
| CCB1567 | W168 *amyE::pHM2-spyA* (pl925) | | | ^30^ |
| CCB1568 | W168 *rae1::pMUTIN ery amyE::pHM2-spyA* (pl925) | | | ^30^ |
| CCB1572 | W168 *amyE::pHM2-hbsΔ-spyA(UAG)+18nts* (pl927) | | | This study |
| CCB1573 | W168 *rae1::pMUTIN ery amyE::pHM2-hbsΔ-spyA(UAG) +18 nts* (pl927) | | | This study |
| CCB1577 | W168 *rae1::pMUTIN ery amyE::pHM2-hbsΔ-spyA+18nts* (pl895) | | | This study |
| CCB1588 | W168 *rnjA::spc efp::kan pHM2-fliY* (pl856) | | | This study |
| CCB1594 | W168 *amyE::pHM2-bmrX* (pl936) | | | ^30^ |
| CCB1595 | W168 *rae1::pMUTIN ery amyE::pHM2-bmrX* (pl936) | | | ^30^ |
| CCB1596 | W168 *rae1::pMUTIN ery rnjA::spc efp::kan pHM2-fliY* (pl856) | | | This study |
| **Strain** | **Genotype** | **Reference** | | |
| CCB1601 | W168 *amyE::pHM2-spyA (14-ΔLRM)* (pl937) | | | This study |
| CCB1604 | W168 *amyE::pHM2- spyA(hbs)3’UTR* (pl940) | | | This study |
| CCB1605 | W168 *amyE::pHM2-spyA (UGA)* (pl941) | | | This study |
| CCB1606 | W168 *rae1::pMUTIN ery amyE::pHM2-spyA (14-ΔLRM)* (pl937) | | | This study |
| CCB1609 | W168 *rae1::pMUTIN ery amyE::pHM2-spyA(hbs)3’UTR* (pl940) | | | This study |
| CCB1610 | W168 *rae1::pMUTIN ery amyE::pHM2-spyA (UGA)* (pl941) | | | This study |
| CCB1630 | W168 *amyE::pHM2-spyA (17-FKD)* (pl943) | | | This study |
| CCB1631 | W168 *rae1::pMUTIN ery amyE::pHM2-spyA (17-FKD)* (pl943) | | | This study |
| CCB1632 | W168 *amyE::pHM2-spyA (17-RML)* (pl944) | | | This study |
| CCB1633 | W168 *rae1::pMUTIN ery amyE::pHM2-spyA (17-RML)* (pl944) | | | This study |
| CCB1636 | W168 *amyE::pHM2-spyA (16-ΔL)* (pl946) | | | This study |
| CCB1637 | W168 *rae1::pMUTIN ery amyE::pHM2-spyA (16-ΔL)* (pl946) | | | This study |
| CCB1638 | W168 *amyE::pHM2-spyA (15-ΔLR)* (pl947) | | | This study |
| CCB1639 | W168 *rae1::pMUTIN ery amyE::pHM2-spyA (15-ΔLR)* (pl947) | | | This study |
| CCB1640 | W168 *amyE::pHM2-spyA (14-ΔELL)* (pl948) | | | This study |
| CCB1641 | W168 *rae1::pMUTIN ery amyE::pHM2-spyA (14-ΔELL)* (pl948) | | | This study |
| CCB1672 | W168 *amyE::pHM2-gfpbmrX* (pl1014) | | | This study |
| CCB1673 | W168 *rae1::pMUTIN ery amyE::pHM2-gfpbmrX* (pl1014) | | | This study |
| CCB1683 | W168 *amyE::pHM2-spyA+18nts* (pl973) | | | This study |
| CCB1684 | W168 *rae1::pMUTIN ery amyE::pHM2-spyA+18nts* (pl973) | | | This study |
| CCB1685 | W168 *amyE::pHM2-spyA(UAG)+18nts* (pl974) | | | This study |
| CCB1686 | W168 *rae1::pMUTIN ery amyE::pHM2-spyA(UAG)+18nts* (pl974) | | | This study |
| CCB1688 | W168 *amyE::pHM2-spyA (UAA)* (pl977) | | | This study |
| CCB1689 | W168 *rae1::pMUTIN ery amyE::pHM2-spyA (UAA)* (pl977) | | | This study |
| CCB1734 | W168 *amyE::pHM2-gfpspyA (14-ΔLRM)* (pl991) | | | This study |
| CCB1735 | W168 *rae1::pMUTIN ery amyE::pHM2-gfpspyA (14-ΔLRM)* (pl991) | | | This study |
| CCB1736 | W168 *amyE::pHM2-spyAUAGgfp* (pl995) | | | This study |
| CCB1737 | W168 *rae1::pMUTIN ery amyE::pHM2-spyAUAGgfp* (pl995) | | | This study |
| CCB1754 | W168 *amyE::pHM2-bmrXUGAgfp* (pl1005) | | | This study |
| CCB1755 | W168 *rae1::pMUTIN ery* *amyE::pHM2-bmrXUGAgfp* (pl1005) | | | This study |
| CCB1758 | W168 *amyE::pHM2-gfpspyA (6-ΔMLRMELLMHYT)* (pl1006) | | | This study |
| CCB1759 | W168 *rae1::pMUTIN ery amyE::pHM2-gfpspyA (6-ΔMLRMELLMHYT)*  (pl1006) | | | This study |
| CCB1762 | W168 *amyE::pHM2-spyA-AAmod* (pl1008) | | | This study |
| CCB1763 | W168 *rae1::pMUTIN ery amyE::pHM2-spyA-AAmod* (pl1008) | | | This study |
| CCB1785 | W168 *amyE::pHM2-gfpspyA (12-ΔLRMEL)* (pl1012) | | | This study |
| CCB1786 | W168 *rae1::pMUTIN ery amyE::pHM2-gfpspyA (12-ΔLRMEL)* (pl1012) | | | This study |
| CCB1796 | W168 *amyE::pHM2-gfpspyA (13-ΔLRME)* (pl1017) | | | This study |
| CCB1797 | W168 *rae1::pMUTIN ery amyE::pHM2-gfpspyA (13-ΔLRME)* (pl1017) | | | This study |
| **Strain** | **Genotype** | **Reference** | | |
| CCB1798 | W168 *amyE::pHM2-bmrX (16-ΔKTNTEARAVR)* (pl1018) | | | This study |
| CCB1799 | W168 *rae1::pMUTIN ery amyE::pHM2-bmrX (16-ΔKTNTEARAVR)*  (pl1018) | | | This study |
| CCB1820 | W168 *amyE::pHM2-gfpspyA (14-ΔLRM E5Q)* (pl1022) | | | This study |
| CCB1821 | W168 *rae1::pMUTIN ery amyE::pHM2-gfpspyA (14-ΔLRM E5Q)*  (pl1022) | | | This study |
| CCB1822 | W168 *amyE::pHM2-gfpspyA (14-ΔLRM E5D)* (pl1023) | | | This study |
| CCB1823 | W168 *rae1::pMUTIN ery amyE::pHM2-gfpspyA (14-ΔLRM E5D)*  (pl1023) | | | This study |
| CCB1871 | W168 *amyE::pHM2-ΔfliYPP* (pl922) | | | This study |
| CCB1876 | W168 *rae1::pMUTIN amyE::pHM2-ΔfliYPP* (pl922) | | | This study |
| CCB1914 | W168 amyE::pHM2-*bmrXUAG* (pl1056) | | | This study |
| CCB1915 | W168 *rae1*::pMUTIN ery *amyE*::pHM2-*bmrX(UAG)* (pl1056) | | | This study |
| CCB1916 | W168 *amyE*::pHM2-*bmrX+18nts* (pl1058) | | | This study |
| CCB1917 | W168 *rae1*::pMUTIN ery *amyE*::pHM2- *bmrX+18nts* (pl1058) | | | This study |
| CCB1924 | W168 *amyE*::pHM2-*bmrX(UGA)+18nts* (pl1059) | | | This study |
| CCB1925 | W168 *rae1*::pMUTIN ery *amyE*::*pHM2- bmrX(UGA)+18nts* (pl1059) | | | This study |
| CCB1926 | W168 *amyE*::pHM2-*bmrX-AAmod* (pl1057) | | | This study |
| CCB1927 | W168 *rae1*::pMUTIN ery *amyE*::*pHM2- bmrX-AAmod* (pl1057) | | | This study |
| CCB1934 | W168 *amyE*::pHM2-*gfpspyA (14-ΔLRM E5A)* (pl1063) | | | This study |
| CCB1935 | W168 *amyE*::pHM2- *gfpspyA (14-ΔLRM E5K*) (pl1064) | | | This study |
| CCB1936 | W168 *rae1*::pMUTIN ery *amyE*::pHM2*- gfpspyA (14-ΔLRM E5A)*  (pl1063) | | | This study |
| CCB1937 | W168 *rae1*::pMUTIN ery *amyE*::pHM2- *gfpspyA (14-ΔLRM E5K)*  (pl1064) | | | This study |
| CCB1957 | W168 *amyE*::pHM2-*bmrX (11-ΔKTNTEARAVRAKLDE)* (pl1074) | | | This study |
| CCB1958 | W168 *rae1::pMUTIN ery amyE::pHM2-bmrX*  *(11-ΔKTNTEARAVRAKLDE)* (pl1074) | | | This study |

**Table S3.** Plasmids used in this study and constructs

| **Plasmids** | **Upstream fragment** | **Downstream fragment** | **Overlapping fragment** |
| --- | --- | --- | --- |
| pHM2-*hbsΔ-spyA* pl707 | ^26^ | | |
| pHM2-*hbsΔ-spyA (F+1, stop3->Q)* pl735 | ^26^ | | |
| pHM2-*fliY* pl856 | CC2696/CC2697 - template ADNk SSB1002 | | |
| pPspac-*bmrBXgfp* *f0* pl845 | ^30^ | | |
| pHM2-*gfpspyA* pl869 | ^30^ | | |
| pHM2-*spyAgfpspyA*  pl889 | CC1808/2799  template ADNk SSB1002 | CC2800/572  template pl869 | CC1808/CC572 |
| pHM2-*hbsΔ-spyA+18nts*  pl895 | CC1607/ CC2813  template pl707 | CC2814/CC572  template pl707 | CC1607/CC572 |
| pHM2-*spyAgfp*  pl899 | CC1808/CC2799  template pl889 | CC2800/CC572  template pl845 | CC1808/CC572 |
| pHM2-*bmrXgfp* (pl904) | CC2888/CC572 - template pl845 | | |
| pHM2-*ΔfliYAA*  pl923 | CC2696/CC2923  template pl922 | CC2922/HP10  template pl922 | CC2696/HP10 |
| pHM2-*ΔfliY* pl922 | CC2696/CC2950 - template pl856 | | |
| pHM2-*spyA* pl925 | ^30^ | | |
| pHM2-*hbsΔ-spyA(UAG)+18nts*  pl927 | CC1607/CC2953  template pl707 | CC2954/CC572  template pl707 | CC1607/CC572 |
| pHM2-*bmrX* pl936 | ^30^ | | |
| pHM2-*spyA (14-ΔLRM)*  pl937 | CC2962/CC2993  template pl925 | CC2994/CC572  template pl925 | CC2962/CC572 |
| pHM2-*spyA (6-ΔMLRMELLMHYT)*  pl939 | CC2962/CC2997  template pl925 | CC2998/CC572  template pl925 | CC2962/CC572 |
| pHM2- *spyA(hbs)3’UTR*  pl940 | CC2962/CC2999  template pl925 | CC3000/HP10  template pl925 | CC2962/HP10 |
| pHM2-*spyA (UGA)*  pl941 | CC2962/CC3001  template pl925 | CC3002/CC572  template pl925 | CC2962/CC572 |
| pHM2-*spyA (17-FKV)*  pl943 | CC2962/CC3029  template pl925 | CC3030/CC572  template pl925 | CC2962/CC572 |
| pHM2-*spyA (17-RML)*  pl944 | CC2962/CC3031  template pl925 | CC3030/CC572  template pl925 | CC2962/CC572 |
| pHM2-*spyA (16-ΔL)*  pl946 | CC2962/CC3033  template pl925 | CC3030/CC572  template pl925 | CC2962/CC572 |
| pHM2-*spyA (15-ΔL)*  pl947 | CC2962/CC3034  template pl925 | CC3030/CC572  template pl925 | CC2962/CC572 |
| pHM2-*spyA (14-ΔELL)*  pl948 | CC2962/CC3035  template pl925 | CC3036/CC572  template pl925 | CC2962/CC572 |
| pHM2-*spyA+18nts* (pl973) | CC2962/CC572 - template pl895 | | |
| pHM2-*spyA(UAG)+18nts* (pl974) | CC2962/CC572 - template pl927 | | |
| pHM2-*spyA (UAA)*  pl977 | CC2962/CC3182  template pl925 | CC3183/CC572  template pl925 | CC2962/CC572 |
| pHM2-*gfpspyA* *(14-ΔLRM)*  pl991 | CC2634/CC3219  template pl869 | CC3220/CC572  template pl925 | CC2634/CC572 |
| pHM2-*spyAUAGgfp*  pl995 | CC1787/CC2799  template pl925 | CC3184/CC572  template pl899 | CC1787/CC572 |
| pHM2-*bmrXUGAgfp*  pl1005 | CC1787/CC3274  template pl904 | CC3275/CC572  template pl904 | CC1787/CC572 |
| pHM2-*gfpspyA (6-ΔMLRMELLMHYT)*  pl1006 | CC1787/CC3219  template pl869 | CC3273/CC572  Template pl939 | CC1787/CC572 |
| pHM2-*spyA-AAmod*  pl1008 | CC1787/CC3306  template pl735 | CC3305/CC572  template pl925 | CC1787/CC572 |
| **Plasmids** | **Upstream fragment** | **Downstream fragment** | **Overlapping fragment** |
| pHM2-*gfpspyA (12-ΔLRMEL)*  pl1012 | CC1787/CC3219  template pl869 | CC3314/CC572  template pl925 | CC1787/CC572 |
| pHM2-*gfpbmrX*  pl1014 | CC1787/CC2910  template pl869 | CC2911/CC572  Template pl936 | CC1787/CC572 |
| pHM2-*gfpspyA (13-ΔLRME)*  pl1017 | CC1787/CC3219  template pl869 | CC3329/CC572  template pl925 | CC1787/CC572 |
| pHM2-*bmrX (Δ16-KTNTEARAVR)*  pl1018 | CC1787/CC3332  template pl936 | CC3333/CC572  template pl936 | CC1787/CC572 |
| pHM2-*gfpspyA* *(14-ΔLRM E5Q)*  pl1022 | CC1787/CC3219  template pl869 | CC3366/CC572  template pl925 | CC1787/CC572 |
| pHM2-*gfpspyA (14-ΔLRM E5D)*  pl1023 | CC1787/CC3219  template pl869 | CC3367/CC572  template pl925 | CC1787/CC572 |
| pHM2-*bmrX(UAG)*  pl1056 | CC1787/CC3487  template pl936 | CC3488/HP10  template pl936 | CC1787/HP10 |
| pHM2-*bmrX-AAmod*  pl1057 | CC1787/CC3481  template pl936 | CC3482/HP10  template pl936 | CC1787/HP10 |
| pHM2-*bmrX+18nts*  pl1058 | CC1787/CC3483  template pl936 | CC3484/HP10  template pl936 | CC1787/HP10 |
| pHM2-*bmrX(UGA)+18nts*  pl1059 | CC1787/CC3485  template pl936 | CC3486/HP10  template pl936 | CC1787/HP10 |
| pHM2-*gfpspyA (14-ΔLRM E5A)*  pl1063 | CC1787/CC3219  template pl991 | CC3512/CC572  template pl991 | CC1787/CC572 |
| pHM2-*gfpspyA (14-ΔLRM E5K)*  Pl1064 | CC1787/CC3219  template pl991 | CC3513/CC572  template pl991 | CC1787/CC572 |
| pHM2-*gfpbmrX (11ΔKTNTEARAVRAKLDE)*  pl1074 | CC1787/CC3330  template pl869 | CC3534/CC572  template pl936 | CC1787/CC572 |
